# Pathogenic Morphological Signatures of Perturbations in Mitochondrial-Related Genes Revealed by Pooled Imaging Assay

**DOI:** 10.1101/2023.09.30.560021

**Authors:** Colin Kremitzki, Jason Waligorski, Graham Bachman, Lina Mohammed Ali, John Bramley, Maria Vakaki, Vinay Chandrasekaren, Purva Patel, Dhruv Mather, Paul Hime, Robi Mitra, Jeff Milbrandt, William Buchser

## Abstract

Mutations in mitochondrial-related genes underlie numerous neurodegenerative diseases, yet the significance of most variants remains uncertain concerning disease phenotypes. Several thousand genes have been shown to regulate mitochondria in eukaryotic cells, but which of these genes are necessary for proper mitochondrial dynamics? We investigated the degree of morphological disruptions in mitochondrial gene-silenced cells to understand the magnitude of genetic contribution to properly functioning mitochondria and to identify pathogenic variants. We analyzed 5,835 gRNAs in a high dimensional phenotypic dataset produced by the image-based pooled analysis platform Raft-Seq. Using the MFN2-mutant cell phenotype, we identified several genes, including TMEM11, TIMM8A, and three NADH Ubiquinone proteins, as crucial for normal mitochondrial morphology in human U2OS cells. Additionally, we found several missense and UTR variants within the genes SLC25A19 and ATAD3A as drivers of mitochondrial aggregation. By examining multiple features instead of a single readout, this analysis was powered to detect genes which had morphological ‘signatures’ aligned with MFN2-mutant phenotypes. Reanalysis with anomaly detection revealed other critical genes, including APOOL, MCEE, NIT, PHB, and SLC16A7, which perturb mitochondrial network morphology in a manner divergent from MFN2. These studies offer insights into the molecular basis for mitochondrial dysfunction, setting the stage for new genomic diagnostics and therapeutic discovery.

## Introduction

Mitochondria are complex organelles that play a crucial role in energy production and various cellular processes including cellular growth, differentiation, and cell death ^1–3^. Neurons are particularly sensitive to mitochondrial disruptions, due to their elongated, complex shape and metabolic burden ^4^. The processes of fission, fusion, and mitophagy induce morphological changes in these organelles, broadly defined as mitochondrial dynamics ^5–7^. When these dynamics are disrupted, mitochondrial morphology can change, as seen in pathogenic mutations of the gene Mitofusin2 (MFN2) in Charcot-Marie-Tooth Disease Type 2A (CMT2A) which often produce aggregated mitochondrial morphology ^8–12^. Although numerous studies have been conducted in the realm of mitochondrial dynamics in relation to human disease, it is difficult to establish a connection between organellar phenotype to genotype ^13,14^. Several thousand genes have been shown to regulate mitochondria in eukaryotic cells, but it is unknown which are required for proper mitochondrial dynamics. Also, the degree of cellular disruption when these genes are knocked out is unknown. Understanding the necessity and magnitude of genetic contribution to properly functioning mitochondria could help define pathogenic variants that cause human disease.

In addition to the well-studied gene MFN2, solute carrier family 25 member 19 (SLC25A19) and ATPase family AAA domain containing 3A (ATAD3A) are two genes that are known to cause disease and play a role in mitochondrial function. Both code for proteins located in the inner mitochondrial membrane. SLC25A19 is thiamine pyrophosphate carrier. Mutations that disrupt the gene and related transport pathways, especially thiamine transport, are linked to several conditions such as microcephaly and other CNS malformations, anemias, neuropathy, aciduria, carnitine deficiency, HHH syndrome, early infantile epileptic encephalopathy type 3, aspartate/glutamate deficiency, Fontaine progeroid syndrome, and citrullinemia type II ^15–17^. ATAD3A encodes a mitochondrial inner membrane protein essential for mitochondrial dynamics, nucleoid organization, protein translation, and cholesterol metabolism ^18–20^. Understanding the impact of SLC25A19 and ATAD3A at the variant level could provide mechanistic insights and set the stage for more targeted therapeutics given their relevance in human disease.

Still, MFN2, SLC25A19, and ATAD3A are only 3 of hundreds of genes that have been identified to have mitochondrial implications in disease. A renewed effort in functional genomics to enhance understanding of the intricate biology behind sequencing data has led to the development of genetic CRISPR screens, including stand-alone and combined formats like survival, FACS-based, arrayed, and single-cell, for greater insight ^21^. Image-based screening techniques linking phenotype to genotype by microscopy allow the interrogation of a comprehensive array of perturbations in live cell populations ^22–24^. Similarly, a recent advent has been the introduction of screening techniques designed to investigate the morphologies of specific organelles and their links to human disease ^12,25^. Deep cellular phenotyping techniques have also emerged exploring molecular characterization, cellular morphology, functional single cell analysis, and even cellular interactions ^26–31^. Such techniques signify a critical shift beyond cell survival and toward a deeper exploration of organelle structure and function.

Here, we utilize Raft-Seq to examine the detailed morphology and activity of mitochondria in single human cells. We used the U2OS osteosarcoma cell line for its large mitochondrial network and extensive characterization with MFN2 variants ^12^. By knocking out the endogenous locus for 1,410 known mitochondrial-related genes, we aim to determine those which produce a morphological signature akin to mutations in MFN2, which are known to cause neurodegenerative disease. Although many of these mitochondrial-related genes are well-studied, there continue to be many surprises regarding their impact on mitochondrial dynamics ^32^. Our findings are carefully validated in independent assays where we assess Cas9 cutting efficiency and cellular morphology across thousands of cells. Not only can we measure the relative disruption of mitochondrial morphology in different knockouts, but we can also determine this for specific indel variants along a particular gene. In this study, we leverage Raft-Seq and the irregular morphological signature of MFN2-mutant cells to ascertain mitochondrial-related genes which, once silenced, exhibit similar mitochondrial disruption and the potential for a pathogenic outcome.

## Results

### Evaluating Mitochondrial Phenotypes in 1,410 Mitochondrial-Related Genes using Raft-Seq

Normally, mitochondria are dispersed densely around the perinuclear region and sparsely networked throughout the cytoplasm of human U2OS cells. These cells display a distinct aggregated mitochondrial phenotype when overexpressing a dominant mutant of MFN2 (L76P or R94Q for example), or when edited to contain an endogenous MFN2 variant ^12^. To identify additional genes that perturb mitochondrial morphology and possibly cause neurodegenerative disorders if mutated, we screened 5,835 gRNAs for 1,410 mitochondrial-related genes (**Table S1**) using Raft-Seq, a pooled high content imaging pipeline (**Figure 1a**). These guides were organized into several pools, packaged into lentivirus, and delivered into U2OS doxycycline-inducible (Dox) Cas9 cells at a low multiplicity-of-infection (MOI, 0.1) to ensure one gRNA per cell before raft plating. In the primary screen, 77 raft plates were seeded, stained, and imaged. Cells were stained to both visualize the extent of all mitochondria (using MitoTracker), and their polarization state (using TMRM). Single-cell data points were acquired then filtered (**Figure S1**), resulting in 455,006 healthy and viable cells, each assessed for its mitochondrial signature.

**Figure 1.**
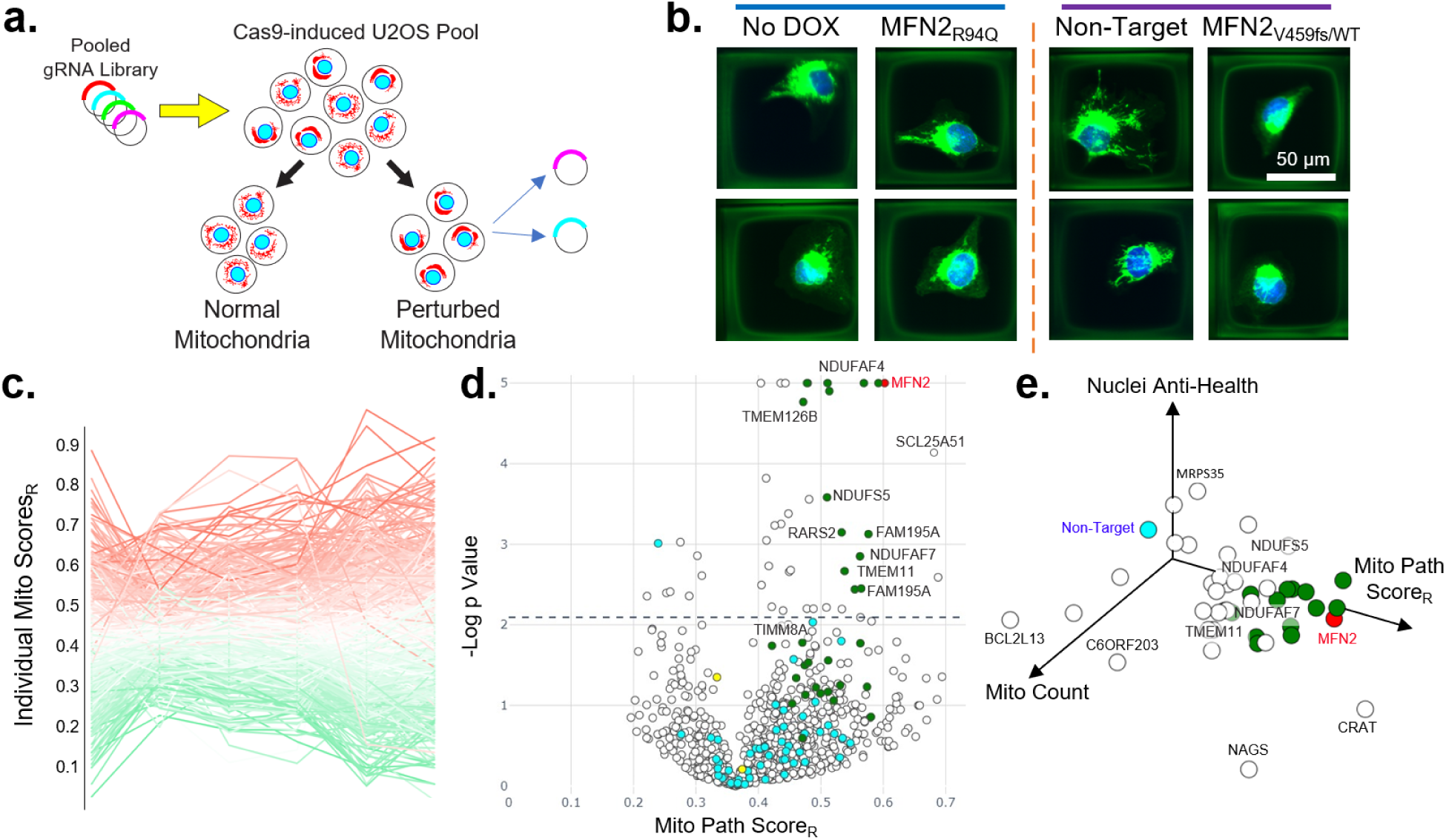
Raft-Seq phenotypic evaluation with mitochondrial-related gRNAs. **a.** Overview of pooled gRNA screening process using Raft-Seq. Mitochondrial-related gRNAs were packaged into lentivirus and infected into U2OS cells with Dox-inducible Cas9. Cells were imaged and analyzed on a subset of phenotypic mitochondrial features before being individually selected for isolation and genotyping. **b.** Fluorescent confocal images of U2OS cells on microrafts to highlight mitochondrial phenotypes of cells from no Cas9 (no Dox, the WT-like control), MFN2_R94Q_ (mutant-like control), Non-Target gRNA (WT-like), and spiked in MFN2_V459fs/WT_ mutants (two examples each). Only the first two (blue line) were ‘labeled’, and used to train the multi-feature classifications, whereas the cells under the purple line were isolated from the mixed screening population and their gRNAs identified through Raft-Seq. **c.** Line plot showing six different multi-feature models to distinguish MFN2 mutant cells from controls. Each line represents the model score for a particular gRNA in the primary screen. Six models are shown as positions along the X-axis, and the connecting points of each line represent the score for a particular gRNA by a particular model. The lines are colored by the average of all the models (Mito Path Score_R_), with red indicating more MFN2-mutant like phenotypes, green control-like, and white intermediate. **d.** Volcano plot listing compiled gRNA results and their position along the X-axis showing WT-like phenotypes (left) or mutant-like phenotypes (right). Blue dots indicate Non-Target gRNAs while green dots represent selections taken into upcoming experiments. Spiked MFN2_V459fs/WT_ positive control indicated with a red marker. - Log P-values of T-tests were capped at 5. Dashed line indicates p-value threshold where FDR is 0.2. **e.** Plot showing gRNA results in relation to Nuclei anti-health (vertical axis), Mito Count (left axis), and Mito Pathogenicity Score (right axis). Nuclei anti-health is calculated as nuclei intensity/nuclei area. Only markers with FDR <= 0.2 are shown.

In the Raft-Seq experiments, the WT-like control consisted of U2OS cells containing the Mito gRNA library where no Cas9 cutting (no Dox) was induced. In contrast, the genetically perturbed control comprised an isogenic line with a mutant endogenous MFN2 (frameshift at V459). These control cell lines were placed in specific wells to serve as anchors, facilitating the distinction between healthy and perturbed in the screen (**Figure 1b**). These controls were used to train ‘vanguard’ logistic regression models to distinguish the MFN2-mutant from the WT-like line. Those model scores were subsequently utilized to select Cas9-active (Dox induced) cells displaying both normal and disrupted phenotypes (**Figure S2**) for Raft-based capture. In total, 29,260 cells were picked into 304 96-well plates and single-cell genotyped to identify the corresponding gRNA. This resulted in 10,730 cells that revealed a single distinct gRNA from the library. In addition, the MFN2_V459fs/WT_ cell line was also introduced to the Dox-induced cells at 0.2% of the total cell count before being plated. Recovery of this “spiked-in” control verified that the Raft-Seq pipeline was successfully able to detect and extract cells with MFN2 mutant-like phenotypes. The result was a highly enriched spiked-in MFN2 control, achieving an AUC of 0.848 (**Figure S3**). Importantly, the gRNA identities weren’t known until after genotyping, and the admixture additionally minimized the impact of batch effects on the score of pathogenicity. Thus, this AUC and enrichment indicate a well-tuned assay.

We trained six multi-feature models consisting of an even mix of binary classifiers (3 Logistic Regressions and 3 Random Forests, 12 random features each) on the normalized data across all plates (**Figure 1c**). Models only saw wild type-like no Cas9 (no Dox) and mutant-like (MFN2_V459fs/WT_) labeled wells for training. Each model used between 9 and 19 mitochondrial features (**Table S2**) to score the differences between the wild type and mutant phenotypes. To prevent bias towards a particular model, we made an ensemble across the six models called the Mito Pathogenicity Score_R_ (“R” indicates raft-based, **Table S3**). Using this Mito Pathogenicity Score_R_, we assembled the gRNAs across multiple observations over well replicates and days. By comparing these scores to the mean of the Non-Target gRNAs, a set of significant MFN2 mutant-like gRNAs emerged (**Figure 1d**). Unlike other types of pooled screens, Raft-Seq can link the gRNA to specific single cells (**Figure S4**). Recovered MFN2_V459fs/WT_ cells were consistently among the most significant results. A subset of the highest scoring gRNAs was selected for further validation.

The Mito Path Score_R_ reduces the dimensionality of the large set of perturbations to a single axis, specifically to detect MFN2 mutant-like morphological signatures. This low-dimensional representation can possibly mask other interesting mitochondrial morphologies (expanded in Underlying Details of the Multi-feature Mitochondrial Pathogenicity Phenotype), as well as phenotypes that may relate more strongly to health or another cellular state (**Figure 1e**). For our downstream analyses, we focused on gRNAs which strongly perturbed the cellular phenotype towards the MFN2-mutant control while not adversely affecting nuclear health as measured by Nuclei Anti-Health (nuclei intensity/nuclei area) ^33^. Such gRNAs result in more clustered mitochondria without detrimental effects on the overall cell health.

### Knockout of TMEM11 and NDU Ubiquinone Proteins Phenocopy MFN2 variants in U2OS cells

To validate the knockout-induced phenotypes elucidated from the screen, we performed an arrayed validation. Two dozen gRNAs were chosen based on a high Mito Path Score_R_, low nuclei anti-health, and low mitochondrial count (**Figure 1e**). We utilized synthetic gRNAs (syngRNAs) (**Table S4**), complexed with Cas9 as ribonucleoprotein complexes which were transfected into U2OS cells. Non-Target, ATP5G3 and PRSS35 gRNA-nucleofected U2OS cells served as the negative controls (the latter two had WT-like results in the primary screen), and a MFN2 gRNA was used as the positive control. These “parent” cell populations were grown out and subsequently split for imaging and cutting analysis. Cas9/syngRNA cutting efficiency was above 95% for most of the syngRNAs (**Figure S5**). Using a plate-based version of the Mito Path Score, the compiled results highlighted a group of syngRNAs which yielded strong MFN2 mutant-like phenotypes (**Figure 2a**, Kruskal-Wallis amongst plate replicates p=3.51×10^-11^). Out of all the validated gRNAs, TMEM11 yielded the highest Mito Path Score_p_, followed by NDUFAF4, TIMM8A, MRPS18C, NDUFAF7, NDUFS5, and RARS2 (**Table S5**). The specific phenotypic signature and the penetrance (fraction of cells that are expressing the mitochondrial disruption) are expected to be different for each of these gRNAs. We observed a highly variable penetrance in the different gRNAs (**Figure S6**) and a consistent aggregated phenotype (**Figure 2b**). We also confirmed the robustness (COD r^2^=0.773) of this phenotype amongst technical replicates performed on different days with different physical experimental layouts (**Figure 2c**). As before, the Mito Path Score_P_ was validated against 296 binary classification models, all of which demonstrated high congruency in the top gRNAs perturbing mitochondrial phenotypes (**Figure 2d**).

**Figure 2.**
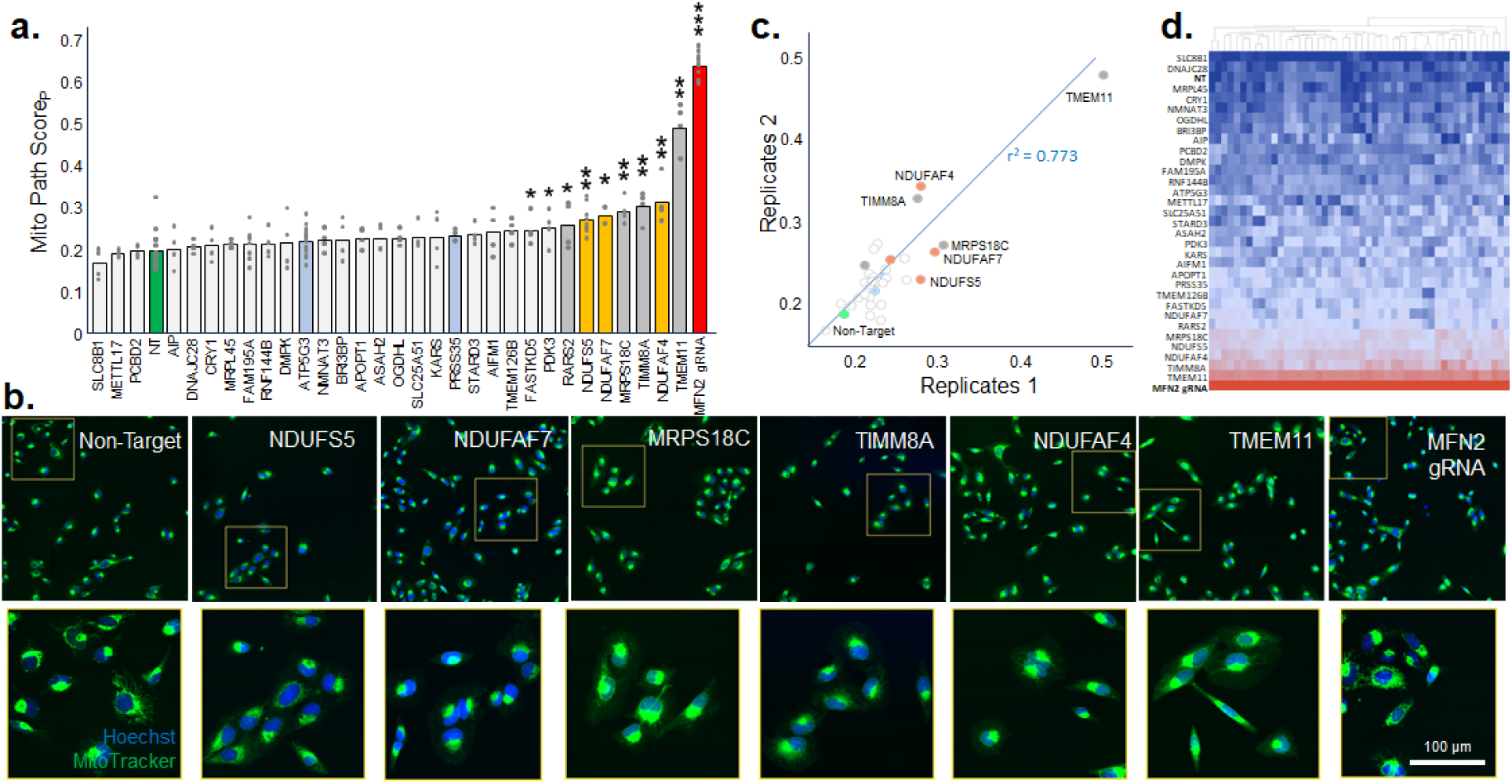
TMEM11 and NDU family members disrupt Mitochondrial Morphology. **a.** U2OS cells nucleofected with syngRNAs representing screen selections and their mitochondrial pathogenicity scores. Significant gRNAs are listed with asterisks denoting P values: * P≤0.05, ** P≤0.01, *** P≤0.001 (Wilcoxon pair test on plate replicates), overall Kruskal-Wallis p = 3.51×10^-11^. Orange bars indicate NADH Ubiquinone (NDU) gene hits, blue bars were purposely selected non-hits, and green and red bars are negative and positive control gRNAs (Non-Target and MFN2 gRNA). Grey bars indicate gRNA selections of significant interest. Markers indicate individual plate replicate scores. **b.** Fluorescent confocal images showing mitochondrial aggregation of a sampling of syngRNA cells and controls. Most of the cells show mitochondrial aggregation similar to MFN2 mutations. **c.** Pearson’s R correlation graph showing positive correlation of replicates for each syngRNA used in this experiment. Mito Path Score_p_ of the replicates populate the axes. MFN2 gRNA was left out of this graph to better visualize the other syngRNAs. **d.** Heat map showing the validation gRNAs and their Mito Path Score_R_ derived from 48 individual binary classification models (blue wild type-like and red MFN2 mutant-like).

Among the genes investigated, TMEM11 had the most penetrant MFN2 mutant-like phenotype. Previously, TMEM11 siRNAs have been shown to reduce mitochondrial tubules, while in Drosophila melanogaster’s TMEM11 ortholog (Pmi) mutations had defects in mitochondrial fission and its phenotype could not be reversed by decreases in OPA1 or MFN2 ^34,35^. Interestingly, recent literature lacks a direct connection of TMEM11 to newly discovered genes that perturb mitochondria. However, TMEM11 has been linked to MTX2, a protein known to be involved in mitochondrial trafficking in neurons (**Figure S7**) ^34^. TMEM11 has also been identified as an outer mitochondrial membrane protein that complexes with BNIP3 and BNIP3L to aid in mitophagy and is required for maintenance of normal mitochondrial morphology ^37^. This is in line with our finding that TMEM11 knockout results in a high Mito Path Score.

It is worth noting that three of the NADH ubiquinone family proteins produced significant Mito Path Score_P_ values. These genes are known to be interconnected, and their network includes TIMM8A and MRPS18C (**Figure S7**). Variants in NDUFAF4/7 are known to cause infantile mitochondrial encephalopathy (MC1DN15) and Leigh Syndrome ^35,36^. TIMM8A variants are associated with Mohr-Tranebjaerg syndrome (deafness-dystonia-optic neuronopathy syndrome), and the gene is part of the mitochondrial protein import system ^37,38^. MRPS18C is part of the mitochondrial ribosomal small subunit and is the least studied of the genes for which knockdown led to disrupted mitochondrial morphology ^39^.

We also examined whether overexpression of *normal* (not mutant) MFN2 could rescue the mutant phenotype caused by TMEM11 or MRPS18C disruption. In addition to regular U2OS WT cells (with endogenous MFN2), we also introduced the syngRNAs into U2OS cells that had been pre-selected for MFN2 overexpression. Neither TMEM11 nor MRPS18C were rescued by the overexpression of MFN2 (**Figure S8**), consistent with previous findings ^40^. Curiously, MFN2 overexpression exacerbated the phenotype of TMEM11 to some extent.

### Scanning Mutagenesis of SLC25A19 and ATAD3A

In addition to gene knockdown, we evaluated the effect of variants in two genes which are relevant to mitochondria and known to cause human disease: the transmembrane protein SLC25A19 and the ATPase ATAD3A. We interrogated a ‘scanning’ gRNA library comprised of 224 gRNAs targeting SLC25A19 (142 gRNAs) and ATAD3A (82 gRNAs) ^41^. Indelphi ^42^ was used to design gRNAs tiled across both genes (**Figure 3a**) with predicted in-frame editing consequences, leading to non-synonymous mutations. Using Raft-Seq, we again undertook pooled image-based analyses of U2OS cells which contained the scanning mutagenesis library. In total, 4,128 cells were captured and genotyped. The isogenic clone MFN2_V459fs/WT_ was again used as the MFN2-mutant control, while the WT-like control consisted of library-containing cells with no Cas9 cutting (no Dox). The Mito Path Score_R_ was used on the primary dataset, and the results revealed positions where gRNAs produced indels that impacted mitochondrial morphology (**Figure 3b**). The gRNA cut sites were also plotted along the cDNA of the respective gene, such that the disrupted domains are highlighted (**Figure S9**), but this revealed no clear areas of cutting enrichment or depletion, likely resulting from frameshift mutations at multiple sites.

**Figure 3.**
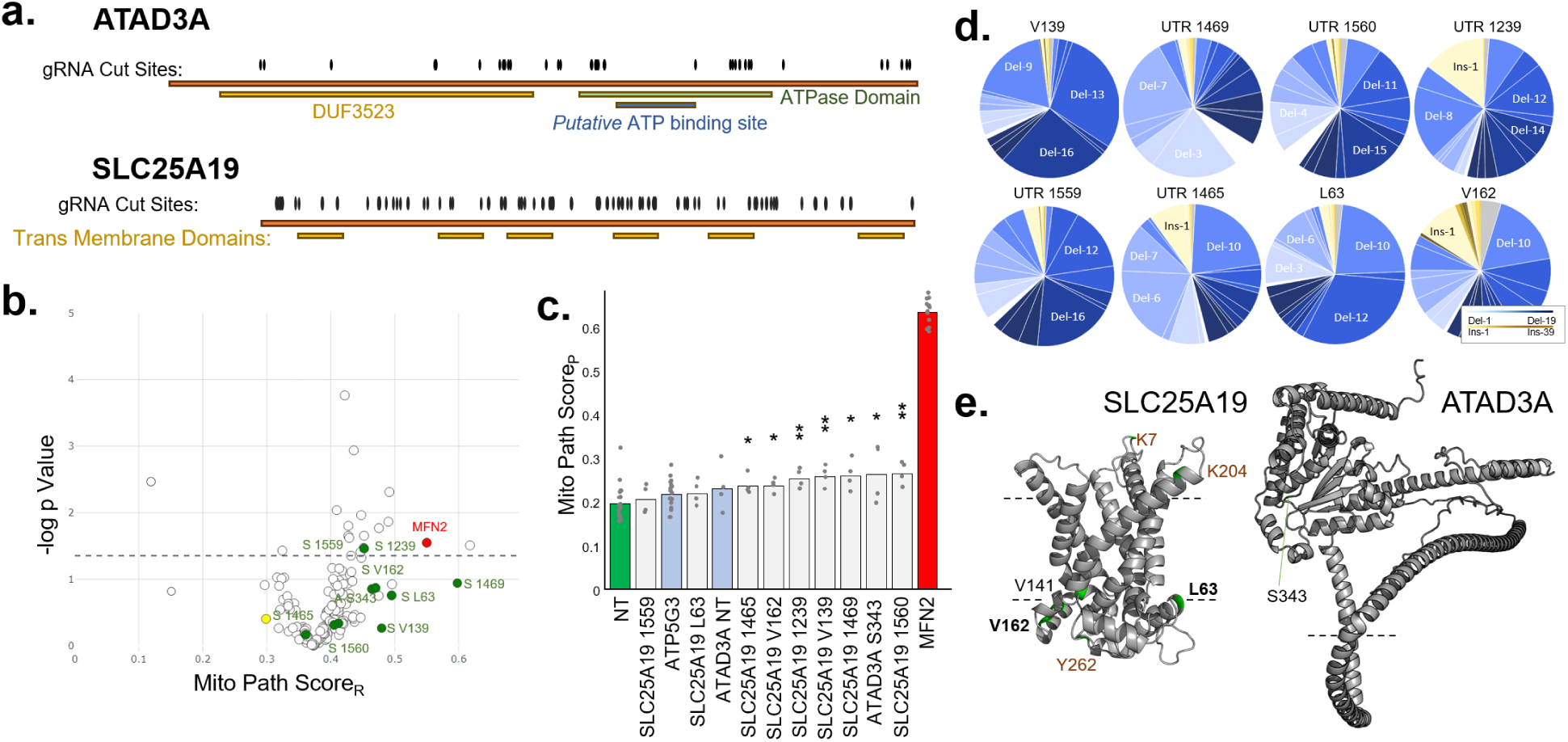
Variants within ATAD3A and SLC25A19 show MFN2-like perturbation of Mitochondrial Morphology. **a.** Diagram of cDNA and domain architecture for 2 genes (SLC25A19 and ATAD3A). gRNA cut sites are shown tiled across each gene from the scanning mutagenesis library. **b.** Volcano plot of results from the scanning mutagenesis library in these two genes. Mito Pathogenicity Score where higher values indicate mitochondrial phenotypes more like MFN2-mutant cells. -log p-value score determined by Mann-Whitney U test against Non-Target gRNAs. Dashed line indicates p-value threshold where FDR is 0.2. **c.** Results from arrayed validation of the top 8 gRNAs identified in the scanning mutagenesis. Genes selected for validation ranked by Mito Path Score_P_. Green denotes non-target, Blue anti-hits, and Red MFN2 gRNA. Overall Kruskal-Wallis p = 4.43×10^-8^, stars indicate significance as *=p≤0.05 and **=p≤0.01 and gray dots show individual replicate scores. **d.** Pie charts showing deletion/insertion representation of the 8 syngRNA bulk samples taken just before phenotyping. **e.** Inner mitochondrial membrane-bound representations of SLC25A19 and ATAD3A. For SLC25A19, the upper and lower dashed lines represent the inner mitochondrial membrane bilayer, with the intermembrane space above and matrix below. For ATAD3A, the intermembrane space is depicted above the dashed line. Key residues for both proteins are labeled.

Initially, we ran a small set of multi-feature analyses (**Figure S10**) on the results from the variant screening and selected ten of the highest-ranked gRNAs that scored consistently across the models (**Figure S11**). The highest scoring gRNAs were then taken into arrayed validation experiments (**Table S6**). Using individual syngRNAs, we attained an average of 92% cutting, and evaluated the Mito Path Score_R_ for the set of SLC25A19 and ATAD3A variants (**Figure 3c**, overall Kruskal-Wallis amongst plate replicates p = 4.43×10^-8^). SLC25A19 syngRNAs, which insert indels near V139 and V162, as well as the 5’ UTR at positions 1465, 1239, 1469, and 1560, all had significantly perturbed mitochondrial phenotypes. Likewise, ATAD3A indels at S343 produced a strongly perturbed phenotype. These results were as expected since those gRNAs had the strongest phenotypes in the microraft-based experiments. Images of the mitochondrial morphology of these variants confirm their disruption (**Figure S12**).

Next, we analyzed the genomic consequences using bulk samples taken from the same cell lines used for the phenotypic analysis. Generally, the gRNAs resulted in in-frame indels in just under half of the cells (as purposely designed by Indelphi). However, the specific insertions and deletions exhibited wide variation (**Figure 3d**). Additionally, we employed AlphaFold2 to generate the proteins’ 3D structures within the mitochondrial membrane, since SLC25A19 and ATAD3A are both mitochondrial membrane proteins ^43^. The high Mito Path Score_P_ mutations tended to fall near the membrane leaflet interface for SLC25A19, and at a single loop in ATAD3A (**Figure 3e**). While these variants did not induce as pronounced a perturbed phenotype as in MFN2, they appeared to have a subtle yet robust effect on mitochondrial morphology, comparable to the NDU knockdowns (**Figure S13** for comparison).

### Underlying Details of the Multi-feature Mitochondrial Pathogenicity Phenotype

In an effort to discover genes essential for normal mitochondrial morphology in U2OS cells, we conducted a comprehensive assessment of the mitochondrial phenotype across several dimensions. To understand the contribution of each dimension to the MFN2 phenotypic signature, we employed the following approach. Each 20x image of a cell was segmented on the Hoechst nucleus, and then a 12 µm region was expanded to account for the rest of the cell body. Pixels were measured separately for MitoTracker and TMRM intensity, producing 66 numeric features. These features were standardized and ranked by the Kolmogorov-Smirnov (KS) statistic comparing the labeled controls, MFN2_V459fs/WT_ versus the WT-like No Cas9 (no Dox) condition (**Figure 4a**). The integrated intensity of the perinuclear mitochondria (Nuclei Total Intensity) emerged as the most discriminative single feature (**Figure 4b**), while texture features (which measure the variation of pixel intensity within a region of the cell) also scored a high KS statistic (**Figure 4c**). However, no single feature adequately separated the labeled controls. Simple linear combinations of pairs of features also did not improve the separation between controls (**Figure 4d**), but a multi-feature machine learning model did (**Figure 4e, Figure S3**). Random forest classifiers demonstrated greater discriminatory power overall compared to Logistic Regressions, but also produced overly optimistic AUC scores when inferring on data seen during training, while Logistic Regression produced more conservative AUC scores under the same conditions.

**Figure 4.**
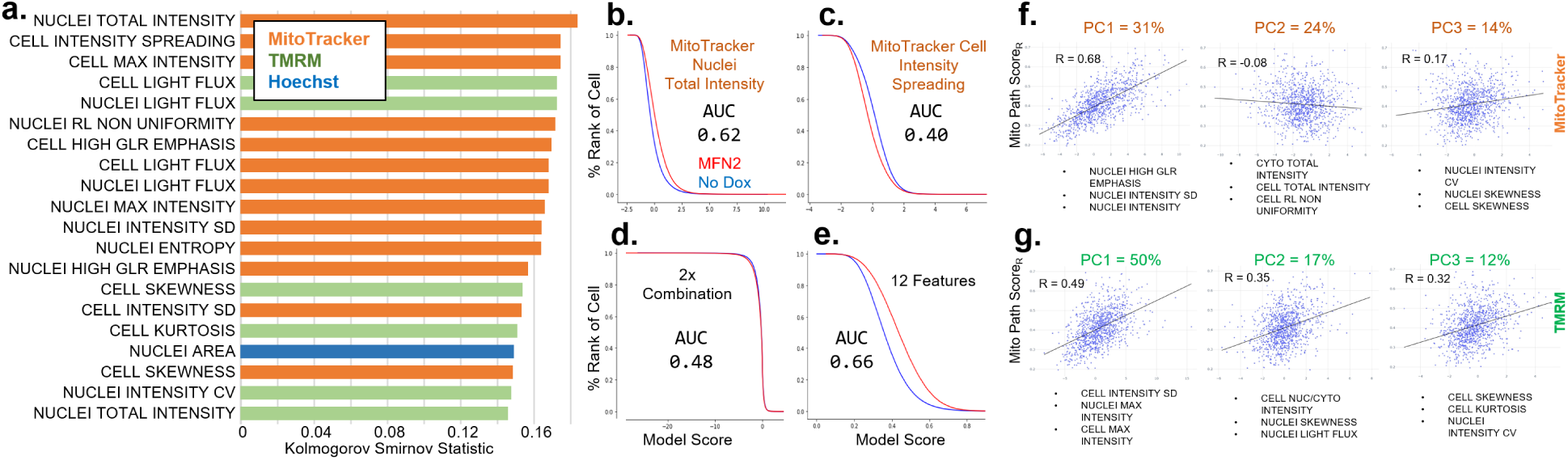
Mitochondrial Phenotype Composition of Individual Features and Principal Components. **a.** Twenty most discriminative features between MFN2_V459fs/WT_ positive controls and No Cas9 (no Dox) negative controls in mitochondrial gene knockout screening data as determined by the Kolmogorov-Smirnov test statistic. Bars are color-coded according to the wavelength from which each feature was derived. Every bar shown had a KS p-value near 0. **b-e** Cumulative probability histograms from the full dataset with red line showing MFN2_V459fs/WT_ cells and blue line No Cas9 cells. The stated AUCs are for the full dataset (training and test). **b.** Cumulative probability histogram segmented according to the most discriminative feature: MitoTracker nuclei total intensity. **c.** The second most discriminative feature: MitoTracker cell intensity spreading. **d.** The product of the top two features. **e.** One of the 12-feature logistic regressions from the Mito Path Score_R_ ensemble. **f.** Scatter plots and the Pearson’s R correlation between the Mito Path Score_R_ and each of the top three principal components for MitoTracker and TMRM. The top loadings for each principal component are included. The percentage listed after the PC1, 2, and 3 are the percent explained variance for that PC.

Next, we independently decomposed our highly dimensional dataset using principal component analysis (PCA) for the MitoTracker and TMRM features (**Figure 4f-g**). Cell-based comparisons against the PCs are also included (**Figure S14**). For MitoTracker, the first 3 PCs explained 69% of the total variance. MitoTracker PC1 happened to strongly correlate (Pearson r = 0.68, p = 2.8e-64) with the Mito Path Score_R_, indicating that the morphological disturbance caused by the MFN2 mutation represented a large fraction in the data overall (31% variance explained), but not all of it. The top 3 feature loadings into PC1 were all perinuclear mitochondrial features that were related to the ‘texture’ or variation among the MitoTracker pixel intensities. The TMRM PCs were also moderately correlated to the Mito Path Score_R_ (Pearson R = 0.49, 0.35, and 0.32), indicating that the variance in TMRM wasn’t as well linked to the MFN2 phenotypic signature – likely because TMRM indicates mitochondrial activity and not morphology. Two of the top three feature loadings for TMRM PC1 were “Max Intensity”, which is the brightness of the strongest mitochondrial pixel in the TMRM wavelength, which could be interpreted as the activity of the cells’ most active mitochondria. All of these analyses were performed on the data spanning the entirety of the microraft-based experiments so encompass a large set of batch variations, making them more generalizable than if taken from a single experiment.

### Unique Mitochondrial Morphologies caused by knockdown of Mitochondrial Genes

Aside from the focus on MFN2-mutant pathology, we sought to identify genes which perturbed mitochondria in distinct ways. Unlike the previous method of scoring phenotypes using classifiers trained on WT and MFN2 mutant cells, here we trained classifiers solely on WT controls—as to perform anomaly detection. This approach enabled us to utilize our existing phenotypic data to identify potentially pathogenic variants whose phenotype is not characterized primarily by the aggregated mitochondria of the MFN2 mutant cells. The combined outputs of several anomaly detection models (Anomaly Score_R_) pinpointed mitochondrial-related genes that, when silenced, resulted in anomalous cell phenotypes compared to cells that were nucleofected with a Non-Target gRNA (**Figure 5a**). Among the most anomalous of these gRNAs were PLGRKT, COX7B2, MRPL39, and AIFM2. Although these gRNAs did not lead to aggregated mitochondria as in MFN2 mutants, they significantly affected mitochondrial morphology (**Figure 5b**, additional images in **Figure S15**).

**Figure 5.**
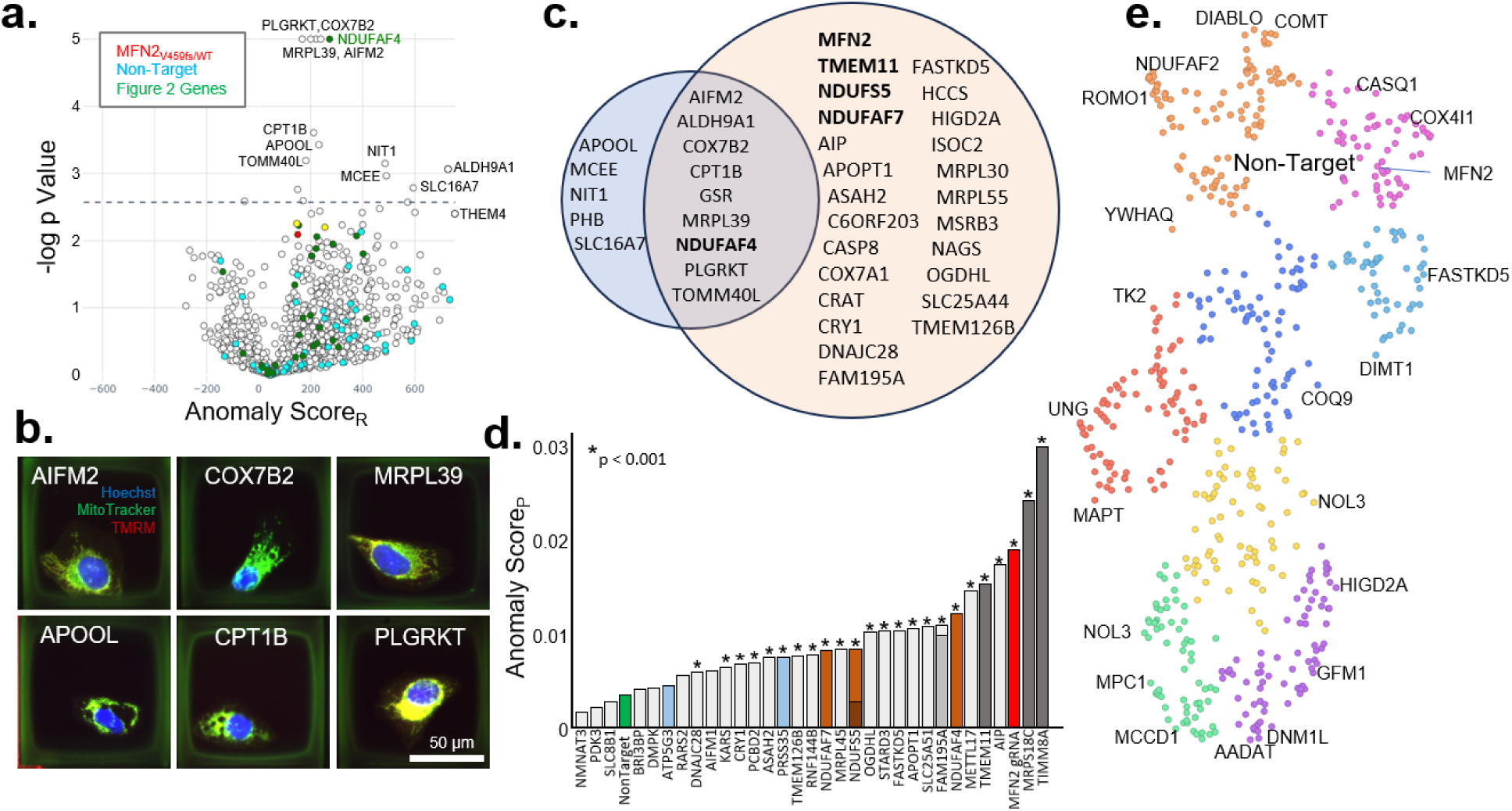
Anomaly Detection to Uncover Additional Mitochondrial Phenotypes Distinct from MFN2. **a.** Volcano plot listing the Anomaly Score_R_ on the x-axis, and the -log p-Value on the y-axis (p-value derived by T-test, with cutoff based on FDR of 0.2). Red is MFN2, Blue represents Non-Targets, Green represents genes that were selected for validation in the primary screen for their MFN2-mutant phenotype, and Yellow represents genes previously selected for their benignity. **b.** Fluorescent confocal images of anomalous single cells on microrafts: Blue=Hoechst, Green=MitoTracker, Yellow=MitoTracker/TMRM colocalization. **c**. Venn Diagram showing genes above the FDR threshold by the Anomaly Score (blue), and the Mito Path Score from Figure 1 (orange). Genes listed in the center are significant for both metrics. Bolded genes had strong Mito Path Score_P_ in the Figure 2 Validation set. **d.** Previously validated gRNAs ranked by their Anomaly Score_P_ instead of their Mito Pathogenicity Score_p_. NDU genes listed as Brown, with NDUFS5 and FAM195A containing two different gRNAs. Asterisks indicate p≤0.001 (p-values derived by Mann-Whitney U-test comparing against Non-Target). **e.** UMAP displaying the position of the single most anomalous gRNA per gene. Non-target gRNAs surround the region indicated, and several genes at the extremes of each region are labeled. Different genes are colored based on hierarchical clustering.

As expected, the Anomaly Score_R_ identified pathogenic phenotypes that were not identified using the Mito Path Score_R_ (**Figure 1**). **Figure 5c** displays the name of the gene identified by either Anomaly Score_R_, Mito Path Score_R_, or both. The overlapping region includes the validated gene NDUFAF4 and eight others that were determined to be significantly pathogenic by both anomaly detection and binary classification models (**Figure 5c**). In addition, five genes were found to be significant only by the Anomaly Score_R_: APOOL, MCEE, NIT1, PHB, SLC16A7. Notably, APOOL has been shown to regulate mitochondrial cristae morphology ^44^. While the link between some of these genes and mitochondrial dynamics is unknown, knocking them out is, in this case, sufficient to affect cell morphology.

Anomaly detection models were also trained on the arrayed data from **Figure 2**, and their combined outputs (Anomaly Score_P_) were used to validate pathogenic phenotypes without relying on the MFN2- mutant signature (**Figure 5d**). According to Anomaly Score_P_, TIMM8A and MRPS18C emerged as the top mitochondrial-anomalous genes, and AIP and METTL17 became more significant than they were according to Mito Path Score_P_.

An alternative to our supervised approaches of binary classification and anomaly detection is the unsupervised algorithm *uniform manifold approximation and projection* (UMAP). We selected one gRNA for each gene based on its maximal difference from the Non-Target gRNA, and then UMAP embedded the gRNAs based on their mitochondrial features (**Figure 5e**). Non-Target gRNAs were grouped together while other subclusters emerged. Examining the outliers within these clusters provides another avenue to identifying potentially interesting morphologies observed in our phenotypic data.

## Discussion

We undertook a pooled image-based phenotype analysis of nearly 6000 gRNAs aimed at perturbing mitochondria. Using high quality single cell images, we assessed mitochondrial morphology (with a general mitochondrial dye) and polarization state (with TMRM). Our analysis was guided by a known genetic mutation in the gene MFN2, which served as a reference to define the abnormal mitochondrial signature, without biases to a particular pre-conceived phenotype like aggregation. The assay worked successfully, with a spiked-in control demonstrating high fidelity enrichment. We found several gRNAs that could significantly perturb mitochondrial morphology along the MFN2-mutant axis and undertook arrayed experiments to validate the effects of the gRNAs independent of the primary raft-based experiment. TMEM11 and TIMM8A were some of the strongest validated genes, as well as a suite of three NADH Ubiquinone proteins NDUFAF4, NDUFAF7, and NDUFS5. In addition to screening pools of mitochondrial gRNAs using Raft-Seq, we also focused in on two specific genes-of-interest that are under-studied yet important to mitochondrial dynamics—SLC25A19 and ATAD3A. We found several sites in the genes that, when mutated, also led to mitochondrial perturbation. Details on the nature of the phenotype that is identifying all these genes involve specific mitochondrial patterns, which are examined in Figure 4. Finally, we re-analyzed the cellular phenotypes to look for anomalous mitochondrial morphologies, extending beyond MFN2-mutant specific morphologies, and uncovered additional genes like APOOL, MCEE, NIT1, PHB, and SLC16A7. Overall, this manuscript offers causal links between gene knockouts and gene-specific variants into the assembly and maintenance of mitochondrial morphology and dynamics.

As a pooled phenotyping platform, Raft-Seq has several strengths which allowed for this dataset and discoveries. Data is retrieved on a per-cell basis, rather than in bulk format, as is the case with enrichment or FACS-based screens ^21,24^. It is most similar to *In Situ* Sequencing, that has been recently pioneered in a genome-wide phenotyping effort ^45^. In that paper, every cell is genotyped after stripping the stains from imaging and doing several rounds of sequencing. In contrast, Raft-Seq only selects cells of interest after imaging, focusing the sequencing resources on a subset of cells. It also means that any type of sequencing can be performed, DNA seq, as is done in this paper to discover the gRNA, but also RNA seq in the style of FACS-Seq, on the captured wells. Raft-Seq enables the selection of live single cells with various indel mutations from the primary phenotyping, and several isogenic clones are available for SLC25A19 and ATAD3A upon request. Additionally, it is worth mentioning the advent of tools in the realm of machine learning-enabled variant prediction, including AlphaMissense and CPT ^46,47^. While such tools excel in predicting substitution variant effects, they face limitations in their ability to predict the functional consequences of indel missense variants. The intricate and subtle nature of such variant effects also poses a challenge for conventional methods like survival or reporter assays, demanding an image-based approach.

Since we are interested in elucidating genes that may lead to mitochondrial-related neuropathy, we focused most of the manuscript on MFN2-mutant like phenotypes. Much work, including our own, has found that MFN2-induced changes in mitochondrial fusion lead to neurodegeneration ^12^. For this reason, we used the MFN2-mutant cells to define the signature of the mitochondrial perturbation we were looking for, which is a combination of a fairly coarse aggregation phenotype, and other more subtle features about mitochondrial morphology and TMRM polarization in each cell. But since the phenotype data is single images linked to a gRNA, many other approaches could be taken. In Figure 5, we used anomaly detection to train classifiers on just the WT-like cells and scored how distant other gRNAs are from this “central” point. This could help define a mitochondrial ‘space’ that represents multiple dimensions but is still grounded in the WT-like state and supervised from there. Although we also showed an unsupervised UMAP projection of the dataset, this manifold isn’t aware that cells can be wild-type or perturbed, so it may miss the directions in mitochondrial ‘space’ that are of interest. The single-cell data is available for brand new interpretations of the phenotype and links with genotype (**Table S3**).

Another technical note is the use of two different scoring systems in the primary microraft-based experiments compared with the syngRNA plate-based experiments. We have both a Mito Path Score_R_ (raft) and _P_ (plate), as well as a similar pair for anomaly detection. There are two reasons for this. First, the raft topology makes it such that the physical size of the cells is slightly different than in a plate. Also, the structure of the raft is slightly concave, so the cells don’t sit as flat as they do on a plate. But more importantly, all the gRNAs recovered from the microrafts were from cells treated with only Dox to induce Cas9, whereas all of the cells in the syngRNA plates were nucleofected with Grna+Cas9 ribonuclear protein complexes. The nucleofection appeared to change the morphology and affected the controls and conditions alike, but made cross-comparisons between raft and plate experiments challenging. For this reason, the two scoring systems were necessary.

The genes found to strongly perturb mitochondrial morphology are fairly unstudied, such as TMEM11, TIMM8A, MRPS18C, and the NADH Ubiquinone genes. Some of them are known to cause disease in humans, and the data presented here could indicate that they might disrupt axons and cause degeneration, as is known to happen with MFN2-neuropathy ^12^. Interestingly, the genes SLC25A51, FAM195A, AIP, and METTL17 were seen to be strong candidates from the microraft-based screening experiments. However, in the validation (plate-based) experiments, their phenotype was weaker (**Figure 2b**). Surprisingly, when their data was re-examined as part of the anomaly detection scoring in **Figure 5d**, they were among the strongest results, with AIP specifically being almost as ‘anomalous’ as MFN2- mutants, and both FAM195A gRNAs showing up strongly as well. SLC25A51 is a mammalian mitochondrial NAD+ transporter, so it may also relate to the NDU findings ^48^. METTL17 and MRPS18C are both involved in mitochondrial ribosomes and in yeast mitochondrial stress response ^49^. AIP is involved in mitochondrial import, as is TIMM8A ^50^.

It is also worth mentioning that several gRNAs showed strong effects that weren’t discussed above. These include APOPT1, FASTKD5, STARD3, OGDHL, that were pulled from the original screening set, but didn’t show as strong signals compared to MFN2-mutant. But, like SLC25A51, they did show significant differences from the non-target gRNAs, and so are another validated set of genes that perturb mitochondrial morphology. Finally, since most gRNAs did validate in at least one of the two methods for testing them (either in a way similar to MFN2 or in a dimension away from non-target), it is worth listing that the majority of the following genes are also likely to perturb mitochondrial morphology but haven’t been independently tested with syngRNAs. They include ALDH9A1, ASAH2, C6ORF203, CASP8, COX7A1, COX7B2, CPT1B, CRAT, CRY1, DNAJC28, GSR, HCCS, HIGD2A, ISOC2, MRPL30, MRPL39, MRPL55, MSRB3, NAGS, PLGRKT, SLC25A44, TOMM40L.

## Methods

### Culture and Transfection

Human osteosarcoma U2OS cell line (U2OS, ATCC HTB-96) with doxycycline-inducible Cas9 ^12^ was cultured in McCoy’s 5A Modified Medium (16600082, Gibco, Gaithersburg, MD, USA) supplemented with 10% tetracycline-free fetal bovine serum (FBS) (PS-FB3, Peak Bio, Pleasanton, CA, USA). Human embryonic kidney (HEK) 293T cells (CRL-11268, ATCC) line was cultured in Dulbecco’s Modified Eagle’s Medium (11965-092, Gibco) supplemented with 10% FBS (26140079, Gibco), 1% Penicillin-Streptomycin (15140122, Gibco) and 1% non-essential amino acids (11140050, Gibco). Both lines were maintained at 37°C, 5% CO_2_ and were monitored daily for overall cell health and confluency. Cells were grown in either T75 or T150 tissue culture flasks and were passaged at 90% confluency. Trypsinization utilized 0.25% Trypsin-EDTA 1x (25200056, Gibco) on the cells for 5 minutes in the 37°C incubator and the cell suspensions were collected and centrifuged for 3 minutes at 1200 RPM. Cells were counted with either a BioRad TC20 automated cell counter or hemacytometer. Cells were passaged at a 1:10 ratio and were always maintained at a cellular representation of (# gRNAs * 1000 * rF), where rF was the representation factor between 2 and 5 for different sub-pools of the library. Passage number was recorded at each passage and cells were not allowed to exceed 10 passage events before freshly thawed cells were introduced. Testing for mycoplasma was performed bi-annually. Experiments in this paper utilized 100×100 µm single and quad reservoir CellRaft Cytosort raft plates (Cell Microsystems Inc, NC, USA). Prior to plating, raft plates were rinsed 3 times with 1xPBS and allowed to incubate at 37°C, 5% CO2 for 3 minutes between each rinse. Plating densities for each were as follows, Quad 48,000 cells total (12,000/quadrant), Single 32,000 cells total. These wells were then brought up in 300-500µl of their respective medias. Post-plating, raft plates were allowed to incubate for 12-14 hours prior to staining/imaging. Isogenic clone MFN2_V459fs/WT_ was spiked into the screen at a frequency of 0.2%.

### Lentivirus Production and Titering

Lentiviruses carrying the gRNA libraries were packaged into 8×10^6^ HEK 293T cells/well in a 6-well plate (TPP92006, MIDSCI, Fenton, MO, USA), using the TransIT Lentivirus transfection reagent (MIR 6600, Mirus Bio, Madison, WI, USA). gRNA libraries were designed from Brunello gRNAs in seven sub-pools with 4-gRNAs per gene and 5% Non-Target (**Table S1**). These gRNAs were assembled into plasmids by Genome Engineering and Stem Cell Center (GESC @ MGI) at Washington University in St. Louis and their representations were confirmed. The gRNA plasmid libraries were combined with VSVg (Plasmid #8454, Addgene, Watertown, MA, USA) and psPAX2 (Plasmid #12260, Addgene) plasmids in a mass ratio of 0.5/0.5/1.0 for a total of 2µg, before being combined with Opti-MEM media (31985070, Gibco) and TransIT reagent according to the manufacturer’s instructions for each well of a 6-well plate. Roughly 200µl of packaging solution was added to each well of a 6-well plate and incubated at 37°C and 5% CO_2_ for 48-60 hours. Media was collected, centrifuged to remove cell debris, and passed through a 0.22µm PES sterile filter (SLGP033RS, EMD Millipore, St. Louis, MO, USA) attached to a 5mL BD Luer-Lok™ Syringe sterile, single use (309646, BD Biosciences, Franklin Lake, NJ, USA). One-third of the total media volume of Lenti-X Concentrator (631232, Takara Bio, Shiga, Japan) was added to the viral media and refrigerated for 30 minutes at 4°C before being centrifuged for 45 minutes at 1500g and 4°C. Supernatant was removed and the concentrated lentivirus was resuspended in 1/10th initial media volume with 1xPBS (10010023, Gibco). Lentivirus was aliquoted and stored at -80°C in 50µl aliquots. Lentivirus was titered by serially infecting U2OS wild type cells with 1/5, 1/25, and 1/125 dilutions with 1µg/µl Polybrene infection/transfection reagent (TR-1003-G, Sigma) for 24 hours. Cells were trypsinized with 0.25% Trypsin (25200056, Gibco) for five minutes in the 37°C incubator and DNA was extracted with in-house extraction buffer to run qPCR for titer determination. IDT PrimeTime Gene Expression Master Mix (1055770, IDT, Coralville, IA, USA) was used for qPCR, along with primers and probes to detect Albumin and Psi, to determine titer against a lentivirus control standard. An ABI 3100 Cycler was used to run qPCR and determine titer.

### CRISPR/Cas9 gRNA Library Infection and Induction

U2OS gRNA library cell lines were created by lentiviral infection at MOI = 0.1, which was performed in T75 flasks. Lentivirus was added to U2OS iCas9 cells along with 10µg/ml Polybrene (TR-1003-G, Sigma, St. Louis, MO, USA), and allowed to infect cells in a 37°C incubator for 24 hours. Lentivirus media was removed after 24 hours and the cells were allowed to rest for another 24 hours before being selected for 7 days with Puromycin at 8µg/ml (73342, STEMCELL Technologies, Cambridge, MA, USA). Surviving cells were expanded, frozen down, and submitted for bulk NGS sequencing to ensure representation of specific guides. STR profiling, to confirm cell type, was performed using NGS-based analysis by the GESC@MGI. Representation of the gRNAs in the cell lines was performed by bulk PCR and NGS analysis to confirm presence of all gRNAs and their rF (representation factor).

### Staining and Microscopy

Cells were stained with the following vital dye suite: Hoechst 33342 (4.05 µM, Hoechst, Thermo Fisher H3570), MitoTracker Deep Red (MitoTracker, 0.5µM, Thermo Fisher M2246), and Tetramethylrhodamine methyl ester (TMRM, 0.1µM, Thermo Fisher I34361) for 30 minutes at 37°C and 5% CO_2_. A master batch of stain was made the day of screening and kept away from light to avoid batch effects. Post-staining, media was removed, and raft plates were rinsed once with media (1mL of media per quadrant or 5mL of media for Single raft plates). Final media volume of raft plates prior to scanning was 2mL (Single) or 500µl/quadrant (Quad). The plates were placed in a Cell Microsystems plate adapter and imaged on Molecular Devices INCell 6500HS Confocal microscope at 20x 0.45 NA utilizing live cell chamber capabilities (5% CO_2_, 37°C temperature). A custom protocol was developed to maximize fields and field of view overlap (12%). Exposure times for Hoechst (405 nm) and TMRM (561 nm) averaged 0.15 seconds while MitoTracker Deep Red (642 nm) averaged 0.05 seconds. Confocality was used in the 405 and 642 nm and plates were scanned in two halves to expedite image tracing, classification, and picking turnaround time.

### Image Analysis and Quality Control

The in-house custom software “FIVTools” was used to perform quality control and calibrations (https://gitlab.com/buchserlab/fivetools). Semi-manual focus checks ensured that out-of-focus fields were excluded from the analysis. Molecular Devices software, INCarta, was then used to trace U2OS cells and extract their nuclear and cellular features, all of which include texture-based features. A network segmentation algorithm in INCarta was used to find mitochondrial puncta within 12µm of the nuclei. Cell-based features from INCarta were joined with image quality annotations and raft position mapping by FIVTools. Quality control post-tracing was done using Tibco’s Spotfire Analyst software. The inclusion criteria for cells taken to downstream analyses were based on nuclear area, nuclear form factor and distance of the cell from the edge of the raft. Furthermore, cellular intensity, and nuclear area and intensity were used to exclude any rafts that had non-cellular debris or dead nuclei (**Figure S1**). Quality checked cells were used in ML-based modeling.

### Identifying Cells of Interest

The filtered cell data table containing all the nuclear, cell body, and mitochondrial features was imported into a Jupyter notebook running Python (version 3.8.13) within Microsoft’s Visual Studio Code application. Cell features are normalized by plate then multiple machine learning models were created with this dataset and were used to select a picklist of raft coordinates containing cells to be captured. This machine learning setup was a binary classification task, where the features computed by the INCARTA tracing program constituted the predictor variables and the experimental condition of each cell was the target variable. Training was done against U2OS cells with the gRNA library but with no Cas9 (no Dox), versus MFN2_V459fs/WT_ cells. Only features that were computed by the INCARTA program were considered for inclusion in the training dataset. Features that were traced in Hoechst or brightfield, global and background features, and most mitochondrial mask features were excluded from the training data (Hoechst and Brightfield don’t include useful information, background and global are near proxies for the position of a cell on the plate which corresponds directly to the target variable, and mitochondrial mask features consist of a small number of pixels so can add noise).

The training data was preprocessed including min-max normalization, synthetic minority oversampling, thresholding for variance within a feature, and thresholding for correlation between features. Moreover, preprocessing also included selecting between two and twenty MitoTracker and TMRM features at random, selecting the k-best features (according to ANOVA F-value), and selecting based on each feature’s importance according to either a Linear Support Vector Machine or a Logistic Regression model, both using L1-type regularization. In addition, models were often hyperparameter tuned. Mitigation of batch effects was achieved through the consistency of stains, image settings, and extraction settings across all plates in a screening experiment. For batch effect mitigation when combining multiple screening experiments, a multi-dataset integrating algorithm called Harmony was used ^51^.

For a given experiment, multiple machine learning models were trained using a randomized 80-20 training-testing split (the number of models varied considerably between experiments). Each model was evaluated according to a variety of metrics including accuracy, precision, recall, F1-score, Matthew’s Correlation Coefficient (MCC), and Area Under the Curve (AUC). However, the most effective evaluation tool was the models’ discriminatory power on admixed labeled cell populations that were hidden during the training. Each model was used to infer a prediction probability for every cell in the original imported data table. Reliable models generated high prediction probabilities for cells labeled “mutant-like,” low for cells labeled “WT-like,” a 50-50 mix of high and low for “Mix50,” and mostly low for the “Cas9”. The foreknowledge of the expected proportion of “WT-like” to “mutant-like” cells in the “Mix50” and “Cas9” groups was a robust indicator of a model’s performance. This evaluation was commonly represented visually in a ranked cumulative probability histogram (see **Figure 1c**). The prediction probabilities were used to rank each cell in an alternating fashion between the most and least likely perturbed cell. The result was several lists of rafts containing cells that were very likely WT-like and pathogenic mutant-like in equal proportion. These picklists of rafts were sent to the CMS AIR system for cellular isolation and extraction.

### Cell Capture and DNA Extraction

Raft capture is detailed previously ^12^. Briefly, semi skirted 96-well PCR plates (1402-9200, USA Scientific, Ocala, FL, USA) were prepped by dispensing 5µl of extraction buffer (molecular grade water with 10mM Tris-HCl (pH 8.0), 2mM EDTA, 200 µg/mL Proteinase K, and 0.2% TritonX-100) into each well using an Integra 12 channel dispenser and the plates are sealed prior to isolations/picking. Once a 96-well PCR plate is isolated, it is placed in a BioRad T100 thermocycler (65°C for 15 min, 95°C for 5 min) to lyse the cell and extract DNA. PCR plates are spun down at 2000 RPMs for 1 min and stored at 4°C prior to downstream NGS applications. Each 96-well plate is manually checked under a dissection microscope to verify successful isolations in each well. Subsequent isolations require disinfection of the PCR wand and loading additional custom picklists with the rafts of interest.

### DNA Amplification and Sequencing

Extracted DNA was amplified using KOD Hot Start Polymerase (71086-4, Millipore Sigma, St. Louis, MO, USA) following manufacturer’s instructions with primers designed in the U6 region of the gRNA CCIV plasmid. Amplification using KOD was optimized for 30 cycles. A nested PCR approach was used where a larger section of the U6 region was amplified and then performed PCR1 with DeepSeq primers (primers with a complimentary Illumina sequence for i7 and i5 sequencing primer attachment) nested inside that larger amplicon ^52^. This allows the samples to bypass a costly cleanup step.

KOD Primers:

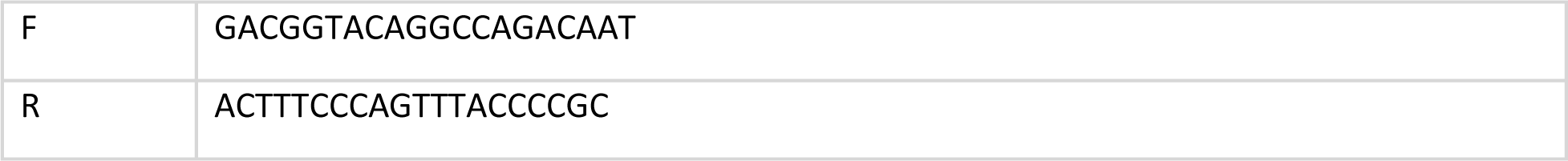

PCR1 DeepSeq Primers:

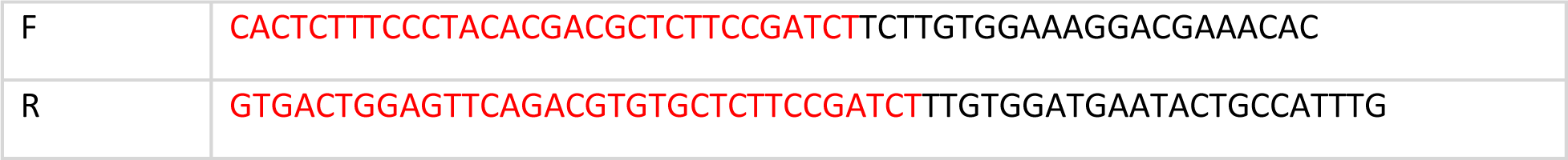

These methods were expanded from a previous paper ^12^. After amplification of the single cell gRNA amplicon, a second amplification was performed to attach a specific 6bp barcoded forward Illumina index primer to the existing forward DeepSeq tags for four 96-well plates with picked single U2OS cells. These primers were used in combination with 4 x 96-well plates of reverse Illumina Index primers from the GESC@MGI to give a unique combination of forward and reverse primers to each well. This allowed the amplified products to be pooled and demultiplexed after next-generation sequencing. A mastermix containing the forward NGS primers, Bioline MyTaq HS Red Mix 2x (C755G97, Merdian Life Sciences, Memphis, TN, USA), and Nuclease Free Water (AM9937, Thermo Scientific, Waltham, MA, USA) was added to 384-well plates (1438-4700, USA Scientific) using a Multidrop Combi Reagent Dispenser (Model 836, Thermo Scientific) and then pooled with 2µL of PCR1 product and reverse indexes using the Biomek I5 liquid handling system (Beckman Coulter, Indianapolis, IN, USA). After pooling of the indexed amplicons on the Biomek, 100µL of pooled sample was cleaned up with 60µl of AMPure XP beads (A63882, Beckman Coulter) according to the manufacturer’s protocol and normalized to the correct concentration using the NanoDrop One Spectrophotometer (Thermo Scientific, ND-ONE-W). Samples were then submitted to the Center for Genome Sciences and Systems Biology (Washington University in St. Louis) for either a 2×250 run or a 2×150 run, depending on amplicon size, performed using the Illumina MiSeq Platform. FastQ files were obtained from demultiplexed Illumina paired reads, then joined, trimmed, and aligned with the various pooled mito gRNA libraries. The resulting dataset showed the number of reads containing each of the 20-mer gRNAs for each well. Typically, a majority of the screened cells were captured after adjusting for isolation and sequencing errors (these filters were applied in Tibco Spotfire), and a flat file as exported containing each picked raft and its assigned genotype.

### Isogenic Cell Line Production

Conditioned McCoy’s media, recovered 1 week after cell culturing to help retain growth factors, was sterile-filtered and 200µl were dispensed across each 96-well round bottom tissue culture plate (TPP 92096) using an Integra 12-channel pipette. Next, 96-well tissue culture plates were allowed to warm prior to isolations. Custom picklists targeted the perturbed cells of interest and were loaded into array records in preparation for isolating/picking. CellRaft AIR System was used to isolate individual rafts into a 96-well flat bottom tissue culture plate. Post-isolation, lids were immediately placed onto the tissue culture plates and were promptly transferred to an incubator at 37°C, 5% CO_2_. Media was added to tissue culture plates weekly (100µl) to account for evaporation. Single cells on rafts were allowed to expand in culture for 2 weeks before they were imaged on Cytiva INCell 6500HS Confocal microscope at 4x in brightfield to evaluate growth. Once wells reached about 40% confluency, they were transferred and progressed through to larger vessels. 10 isogenic clones are available from SLC25A19 and ATAD3A.

### Validation with Synthetic gRNAs

Synthetic gRNAs (syngRNAs) (**Table S4, S6**) from genes of interest were ordered from Integrated DNA Technologies (IDT, Coralville, IA, USA). Primers complementary to 150 bp in either direction of the gRNA were designed and ordered through IDT. U2OS cells were grown in either T75 or T150 tissue culture flasks and were passaged at 90% confluency. RNP complexes consisting of 1µl of syngRNA at 100µM complexed with 10µg of Cas9 protein and stored at -20C the day prior to nucleofection. Nucleofections were performed using 4D-Nucleofector Core and X unit (AAF-1003B, AAF-1003X, Lonza, Morrristown, NJ, USA). In total, approximately 75,000 U2OS cells were nucleofected into small cuvettes (V4SP-3096, Lonza) with 20µl of P3 Primary Cell line solution (V4SP-3096, Lonza) and RNP using the cell type program DS-150. Post-nucleofection, contents from the nucleofection were transferred to 12-well plates (TPP 92012, MIDSCI, Fenton, MO, USA) with 2mL of McCoy’s media and were allowed to proliferate for 7 days, with media changes on day 2 and day 5. The syngRNA cell conditions were randomly arrayed across two 96-well plates using custom software and plated using the Biomek I5 liquid handling robot. After plating, bulk samples were taken from remaining cell suspensions and genomic DNA extractions were performed for downstream NGS/cutting analysis. For the in/del analysis, next generation sequencing reads for each bulk sample were aligned against SLC25A19 and ATAD3A reference sequence to catalogue causative deletions/insertions for each gRNA.

The 96-well plates were incubated at 37°C and 5% CO_2_ overnight. The 96-well plates were stained with 4.05 µM Hoechst 33342, 0.5 µM MitoTracker Deep Red, and 0.1 µM TMRM for 30 minutes at 37°C and 5% CO_2_ as before. A master batch of stain was made the day of screening and kept away from light to avoid batch effects. The 96-well plates were imaged on Molecular Devices INCell 6500HS Confocal microscope at 20x 0.45 NA utilizing live cell chamber. Images were analyzed with INCARTA as before, filtered and compiled. Then, in a Jupyter notebook, multi-feature models were again trained, but using small aNNs (artificial neural networks with 2-3 dense layers) to distinguish between the Non-Target gRNA control, and the MFN2 gRNA control (**Figure 2d, Table S2**).

### Protein Structure

Three-dimensional structures for ATAD3A and SLC25A19 were estimated with the monomer model of AlphaFold2 using the full DBS sequence database ^43^. The highest-ranked model for each protein was selected for transmembrane orientation prediction using the PPM 3.0 web server ^53^. ATAD3A and SLC25A19 both associated with the inner mitochondrial membrane and the N-termini of both proteins were constrained to reside within the transmembrane space ^17,54^. Membrane-bound proteins and specific residues with significant Mito Path Score_R_ were visualized in the PyMOL Molecular Graphis System v2.5.5 (Schrödinger, LLC).

## Definitions

aNN: Artificial Neural Network
AUC: The area under the ROC curve of sensitivity/specificity
Dox: Doxycycline used to induce Cas9 cutting
gRNA: guide RNA for CRISPR Cas9
MFN2: Mitofusin2, a gene that regulates mitochondrial fusion
MFN2-mutant like: Cells with morphological features similar to MFN2-disrupted cells, specifically MFN2_V459fs/WT_
Non-Target gRNA: a scrambled sequence known to not target any human gene
MitoTracker: Vital stain that marks all mitochondria
TMRM: Tetramethylrhodamine methyl ester, a red-orange vital stain that is readily sequestered by active (depolarized) mitochondria
syngRNAs: synthetic gRNA used with Cas9 protein as an RNP
U2OS: Osteosarcoma sarcoma cell line, usually with a dox-inducible Cas9
WT-like: Cells with morphological features similar to wild-type cells. Usually No Cas9 (no doxycycline induction)

## Addenda

## Author Contributions

C.L.K., J.E.W., L.K.MA, G.W.B. and W.J.B. wrote the manuscript. W.J.B., J.C.B., J.E.W., and C.L.K. designed the experiments. Initial planning and ideas brought by W.J.B., J.D.M., R.D.M., J.E.W., and C.L.K.

C.L.K., J.E.W., G.W.B., L.K.MA, J.C.B., M.A.V., P.P. performed the experiments. P.M.H and D.P.M. performed protein structure analysis. J.E.W., C.L.K., G.W.B., L.K.MA, P.P., D.P.M. and W.J.B. analyzed the results. W.J.B., V.D.C., G.B. wrote some of the key software. All reviewed the paper and gave suggestions.

## Acknowledgements

We would like to thank Mariel Liebeskind, Alex Yenkin, Eileen Xu, Samah Nour, Kylan Kelley, Mallory Wright, Serena Elia, Diana Grigore, Josh Milbrandt and Josh Langmade for their help with aspects of the project. Jared Zane and Uma Kaushik helped design some of the primers. We would also like to thank the Milbrandt, DiAntonio and Mitra lab for their continued support. We would like to thank the **McDonnell Genome Institute**, specifically GESC@MGI helped with CRISPR advice and libraries, GTAC@MGI helped with sequencing, as well as WashU’s CGS. Cell Microsystems was very supportive and quick to lend a hand as needed. Finally, we would like to thank the Genetics and MGI administration and our maintenance and cleaning staff.

## Code Availability

https://gitlab.com/buchserlab/fivetools

## Supplemental Tables

**Table S1 gRNA tables with Sequences**

gRNA library sequences and subpools.

**Table S2 Model Features**

Lists the composition of all the relevant machine learning models.

**Table S3 Primary Screening Data (Cell/gRNA based)**

Includes Raft based INCarta data for KO, Scanning Mutagenesis, with Mito Path and Anomaly Detection Scores.

**Table S4 Synthetic gRNA Sequences and Primers**

List of gRNA seqences and the genotyping primers used.

**Table S5 Validation Results (Cell Based)**

Include INCarta data KO, Scanning, with Mito Path and Anomaly Detection Scores.

**Table S6 SLC25A19 and ATAD3A gRNA sequences, synGRNA sequences, primers**

All related to Figure 3 experiments.

