## Supplemental Figures for "Pathogenic Morphological Signatures of Perturbations in Mitochondrial-Related Genes Revealed by Pooled Imaging Assay"

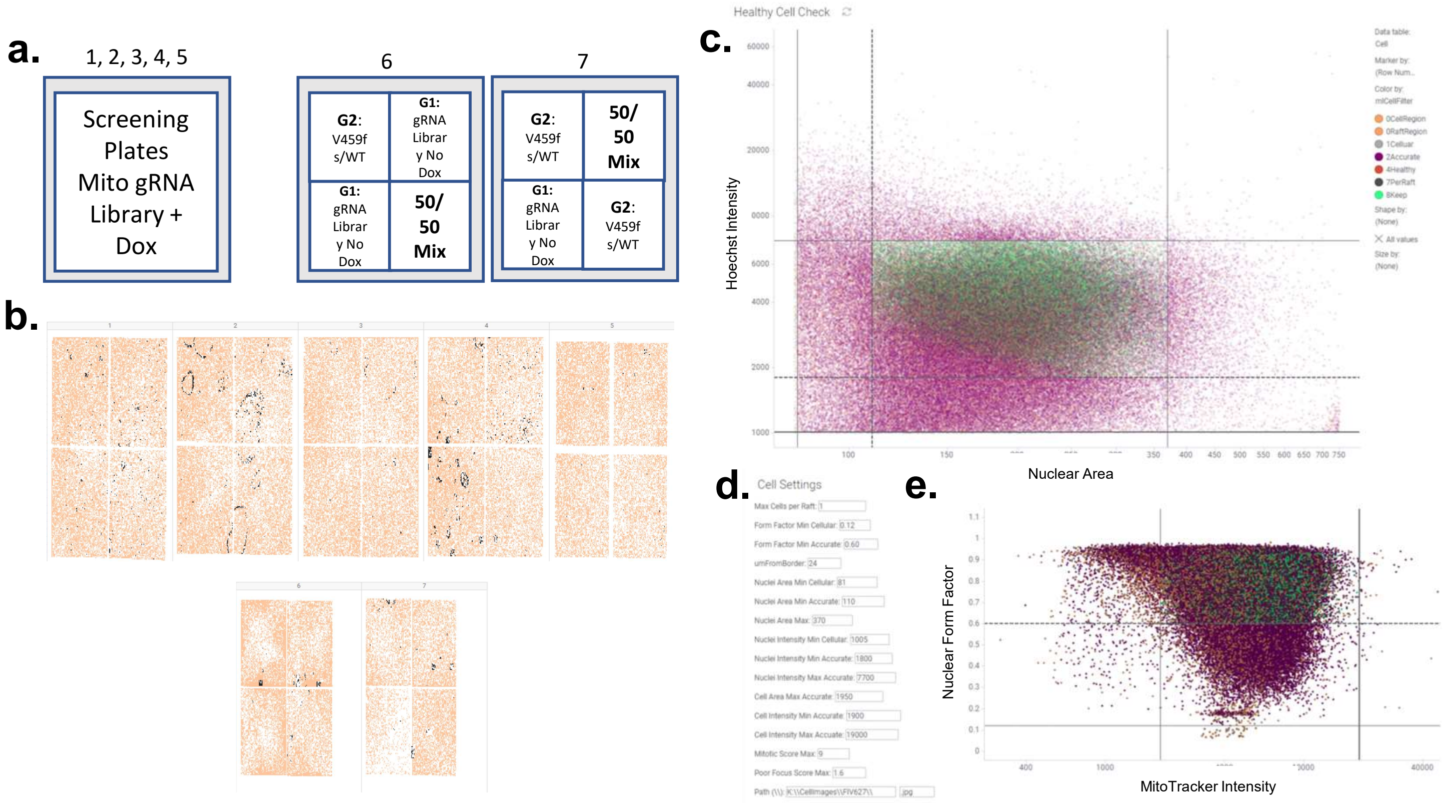


**Figure S1. Raft-Seq Cell Plating and Filtering. a.** Example raft plate layout used for screening gRNA library. Microraft plates 1-5 contain the mito gRNA library + Dox and raft plates 6,7 contain 'labeled' conditions used for anchoring and training machine learning to classify/identify perturbed single cells. **b.** Cell distribution across raft plates during cell-based filtering steps. Orange markers represent microrafts that contain one or more healthy cells, while black markers denote microrafts with cells that were removed during the visual inspection (usually due to poor focus or artifact). White areas are microrafts that do not contain any cells. **c**. Scatter plot where each marker is a unique cell, with nuclear (Hoechst) intensity plotted against nuclear area. Green markers indicate healthy cells kept during the filtering process and purple indicates cells which were omitted. **d**. Adjustable cell-based filtering settings to further define gates for nuclear and cellular area, in addition to focus, and mitotic score prior to downstream analysis. **e**. Scatter plot of individual cells with Nuclei Form Factor (circularity) plotted against MitoTracker Intensity. Cells with Form Factor > 0.6 and MitoTracker Intensity within defined boundaries are kept.

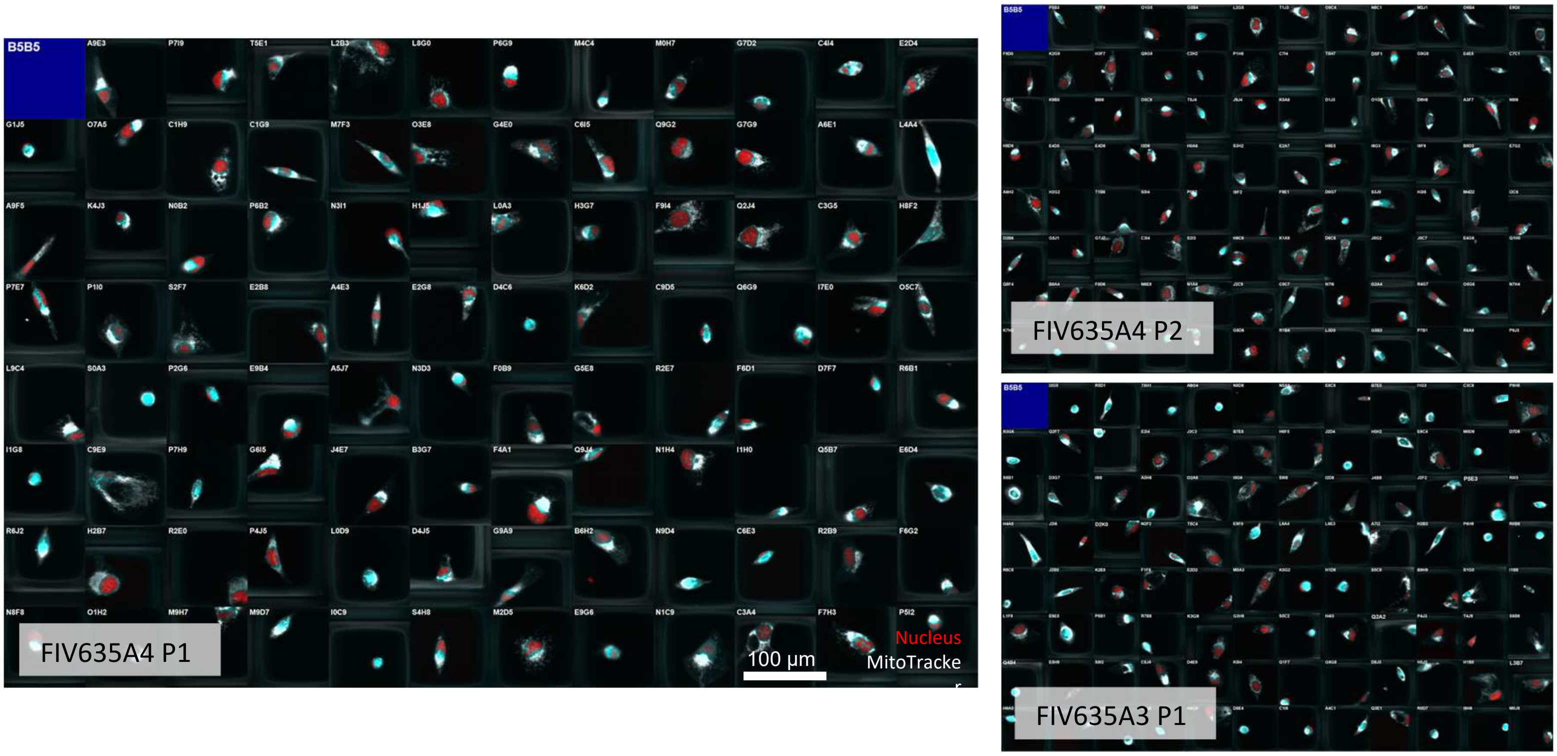


**Figure S2. Confocal Fluorescent Images of Single-Cell Microrafts in Picked Plates.** 20x images of microrafts picked from a raft plate directly into a 96-well plate. Nuclei are visualized in red, mitochondria in white, and regions of overlap are in blue. These microrafts were identified as having cells of phenotypic interest and have been manually inspected to exclude microrafts harboring non-single, dividing, and non-viable cells before sending to the CMS AIR system to be picked. In the upper-left corner of each microraft is a four-character microraft identifier. The blue box is an empty well that was not picked. In the bottom left-corner is a 96-well plate identifier.

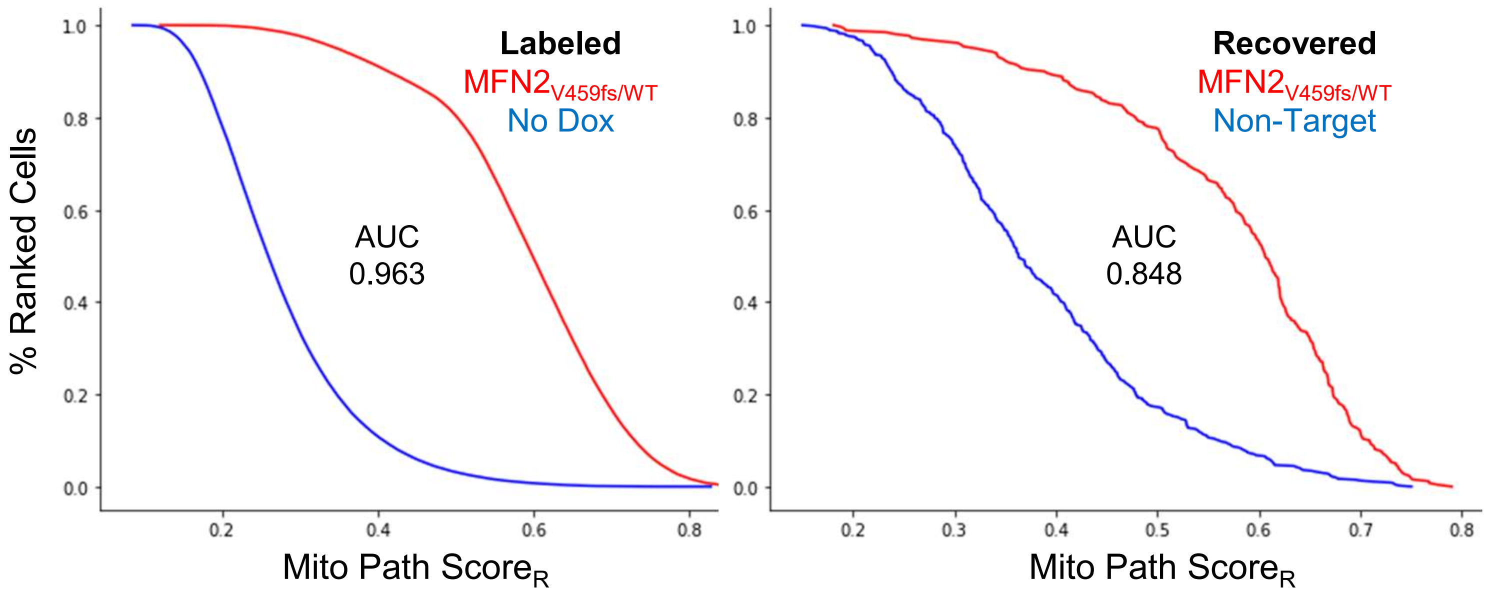


**Figure S3. Recovery of Spiked in MFN2 Mutant Amongst the gRNA Library on Microrafts.** $MFN2_{V459fs/WT}$ was added to the screening wells at 0.2%, approximately the same frequency as a single gRNA. Both graphs are cumulative probability histograms, with the Mito Path $Score_R$ on the x axis, and the % rank of cells which had at least the indicated score on the y-axis. The left graph shows the training data, the 'labeled' wells with known controls. The right graph shows the scores of the recovered gRNAs from within the screening wells. The gRNA identity within the screening (unlabeled) wells was unknown until after microraft selection and single cell genotyping, making for a robust assay evaluation.

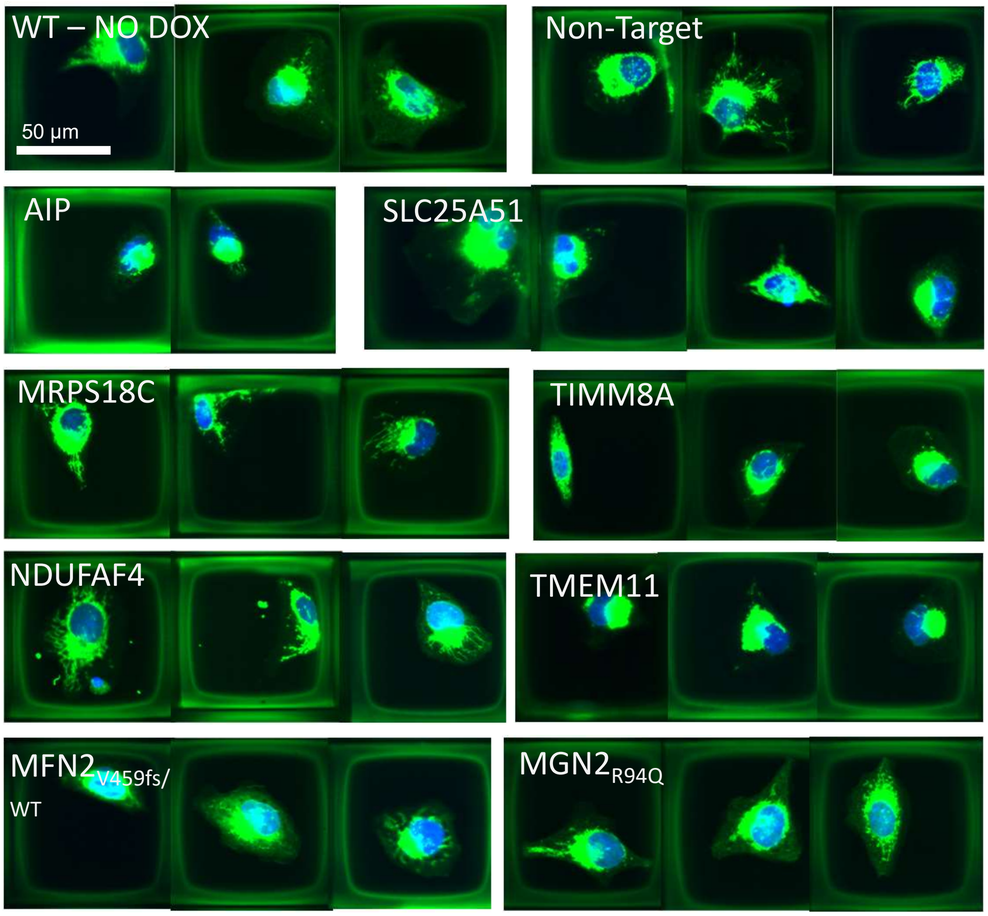


**Figure S4. Examples of Mitochondrial Morphologies in Genotypes of Interest.** Raft-Seq revealed varying mitochondrial morphologies in the genotypes of interest, showcased in these 20x images of U2OS cells on microrafts. WT No Dox (no Cas9) serves as a WT-like control. Similarly, Non-Target gRNAs that don't cut the genome were present in the library, yielding healthy mitochondrial networks. $MFN2_{R94Q}$ and $MFN2_{V459fs/WT}$ are both MFN2-targeting and demonstrated mitochondrial aggregation and disruption, a perturbation resulting from dysregulated fusion/fission. While most genotypes showed varying morphologies, some of the selected gRNAs (listed here) showed strong morphological similarity to the MFN2 mutants. Blue = Hoechst, Green = MitoTracker.

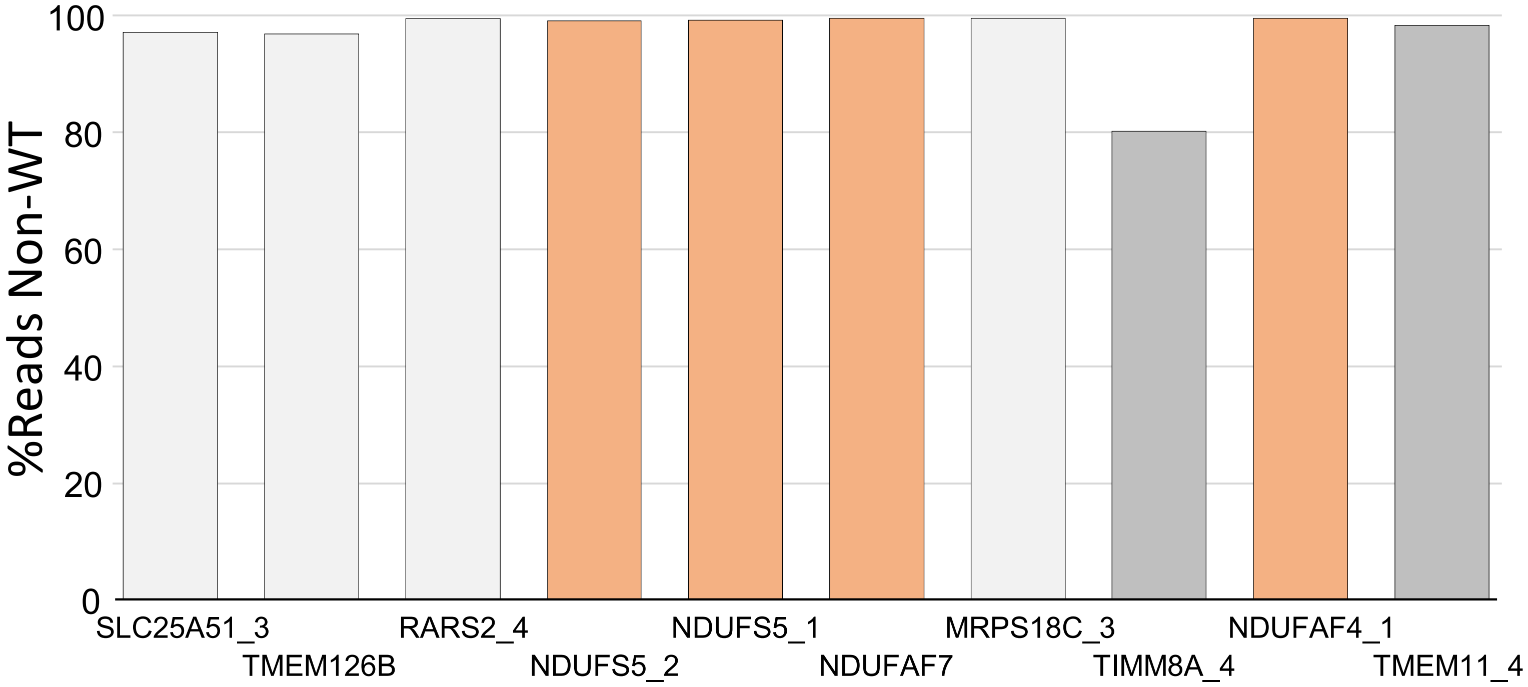


**Figure S5. Cutting Efficiency in Synthetic gRNAs Against Genes-of-Interest.** Bar chart showing CRISPR Cas9 cutting efficiency in the population of cells split off to use for **Figure 2** phenotyping. Efficiency was determined by searching NGS reads for the presence of intact gRNA sequence. If the gRNA 20-mer was not found intact, but the upstream sequences were present, the read was evaluated as non-WT. Orange bars highlight the NDU genes, and the darker grey bars highlight TIMM8A and TMEM11.

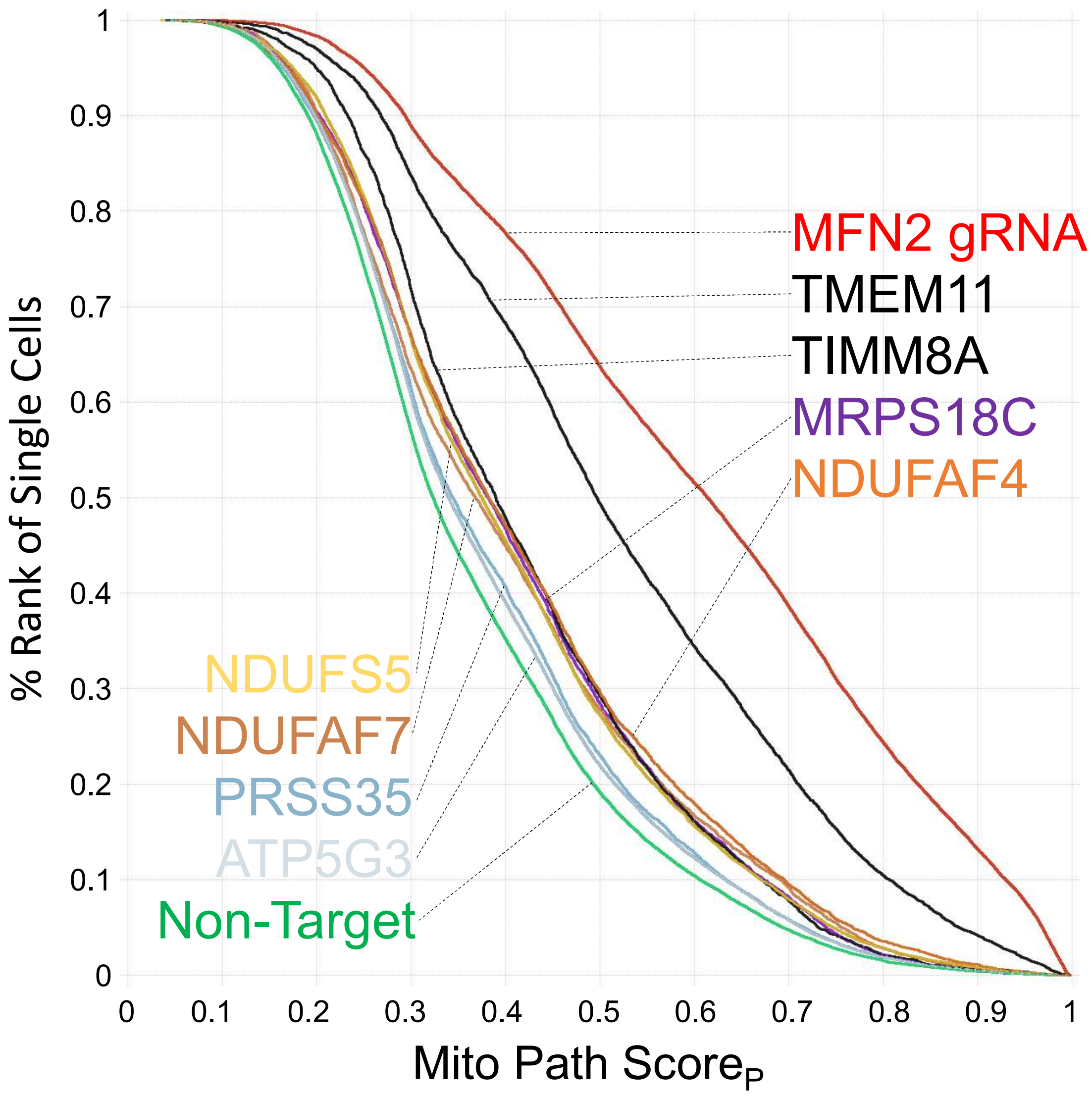


**Figure S6. Penetrance and Variation of Phenotype in Mitochondrial-Related Genes.** Cumulative histogram of individual cells with gRNAs to the indicated genes. X-axis is the Mito Pathogenicity Score shown in **Figure 2**, where higher numbers are more like MFN2 mutations, and lower numbers are more like WT. The Y-axis then ranks each individual cell, so that cells which have more WT phenotypes are shown high, and cells with mutant phenotypes are shown low on the axis. Individual lines are colored by the gRNA they received.

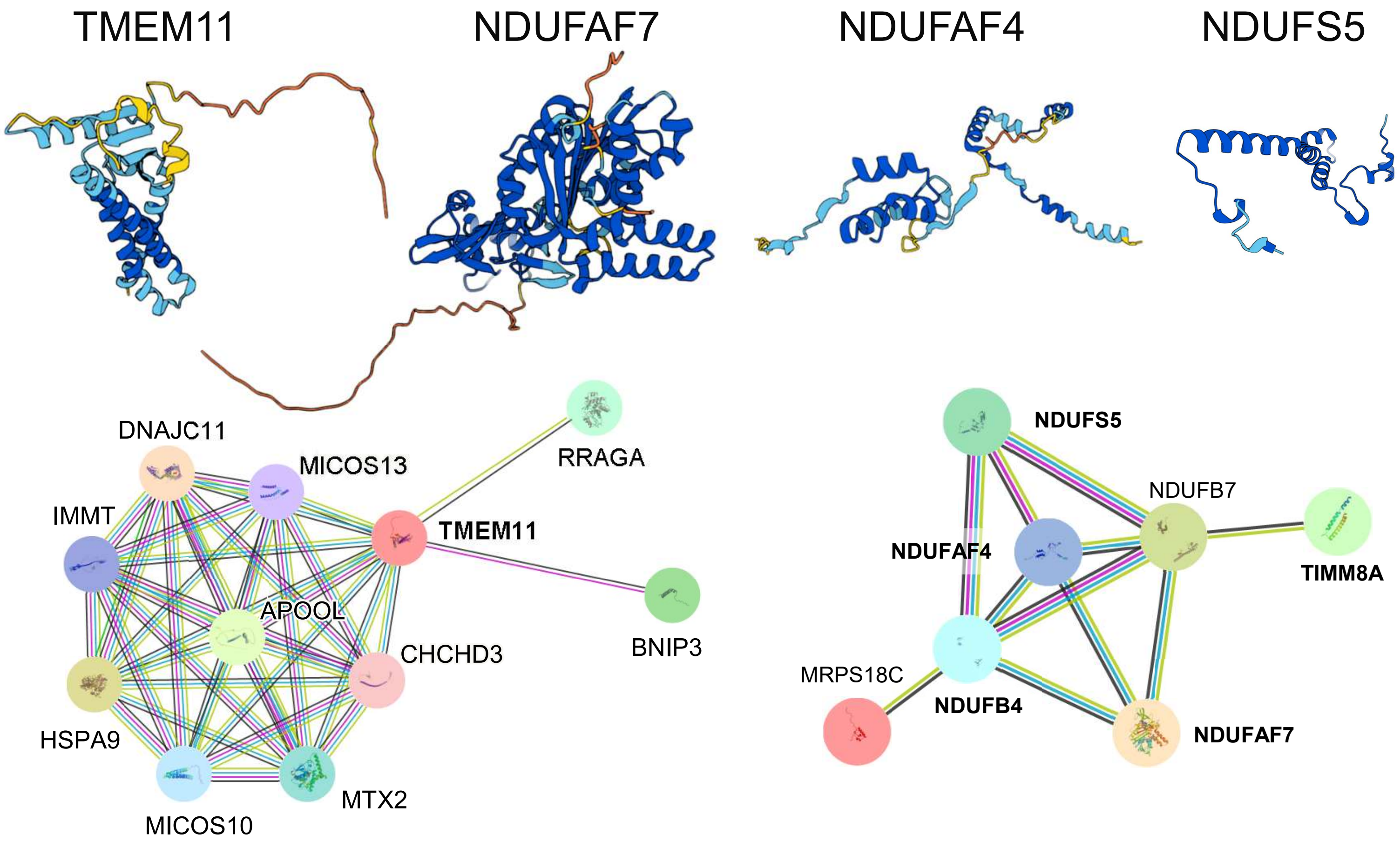


**Figure S7. TMEM11 and NDU Structure and Known Relationships.** Alphafold crystal structures for the proteins-of-interest, where blue indicates high confidence and warmer colors indicate lower folding confidence (from https://alphafold.ebi.ac.uk/). Lower line shows connectivity graph from (https://string-db.org/). TMEM11 is not connected to the other novel genes, but the NDUs are connected to TIMM8A and MRPS18C through two additional NDUs.

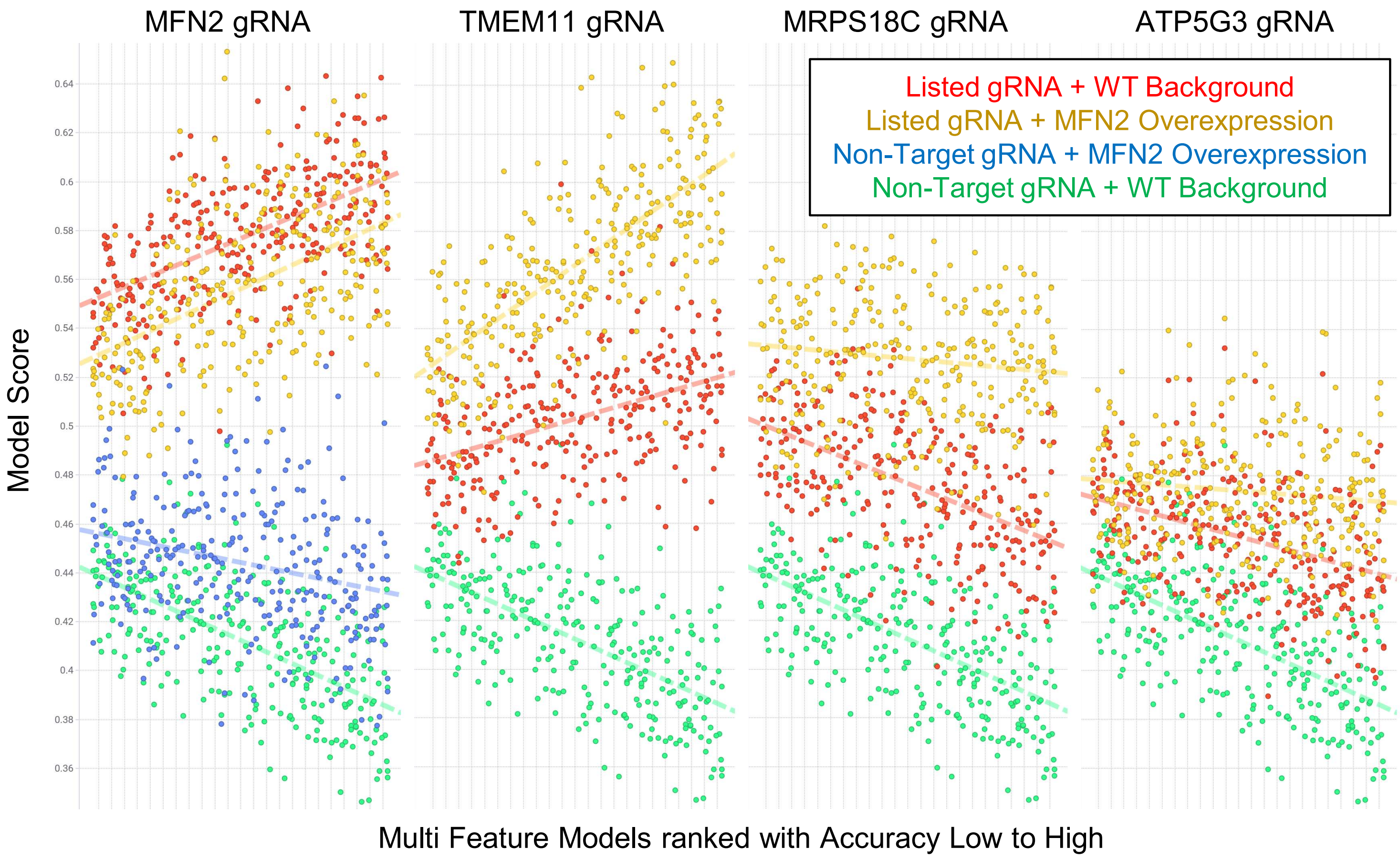


**Figure S8. MFN2 Overexpression Fails to Rescue TMEM11 and MRPS18C Mitochondrial Morphology.** RNPs with the listed gRNAs were transfected into U2OS cells in a WT background or with a stable (normal) MFN2 overexpression. A hundred multi-feature models were constructed with random sets of MitoTracker and TMRM features. These are ordered on the x-axis by their accuracy on discriminating Non-Target from MFN2 gRNA cells in the WT background (accuracies range from 61% to 68%). The prediction score for each model is plotted on the y-axis, where higher number means phenotypically similar to MFN2-mutant cells. Each subplot has the Non-Target + WT background for reference, in addition to indicating a different pair of gRNAs from the screen with or without the MFN2 overexpression.

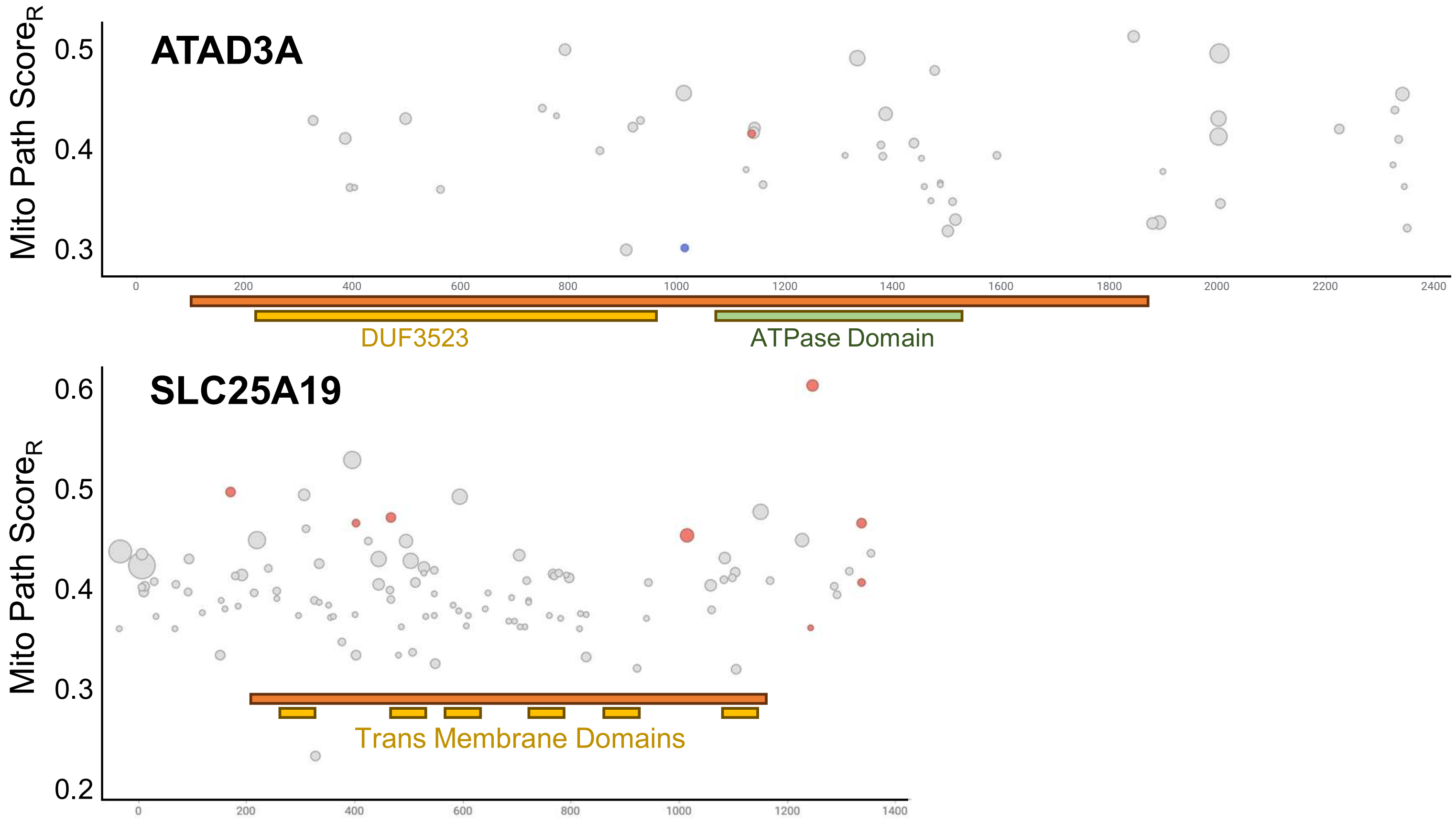


**Figure S9. SLC25A19 and ATAD3A Variant Effects in cDNA Context.** Results from the primary experiment with variants in ATAD3A and SLC25A19. Each marker is a single gRNA which is positioned along the cDNA of the gene listed on the x-axis. The y-axis shows the strength of the phenotype score where higher values are more like MFN2-mutant cells. The size of the markers reflect the p-value from the volcano plot in **Figure 3b**. Red markers were selected for further validation, and the blue marker was selected as a negative control. The protein region of the cDNA is marked with an orange bar and the domain architecture is marked below.

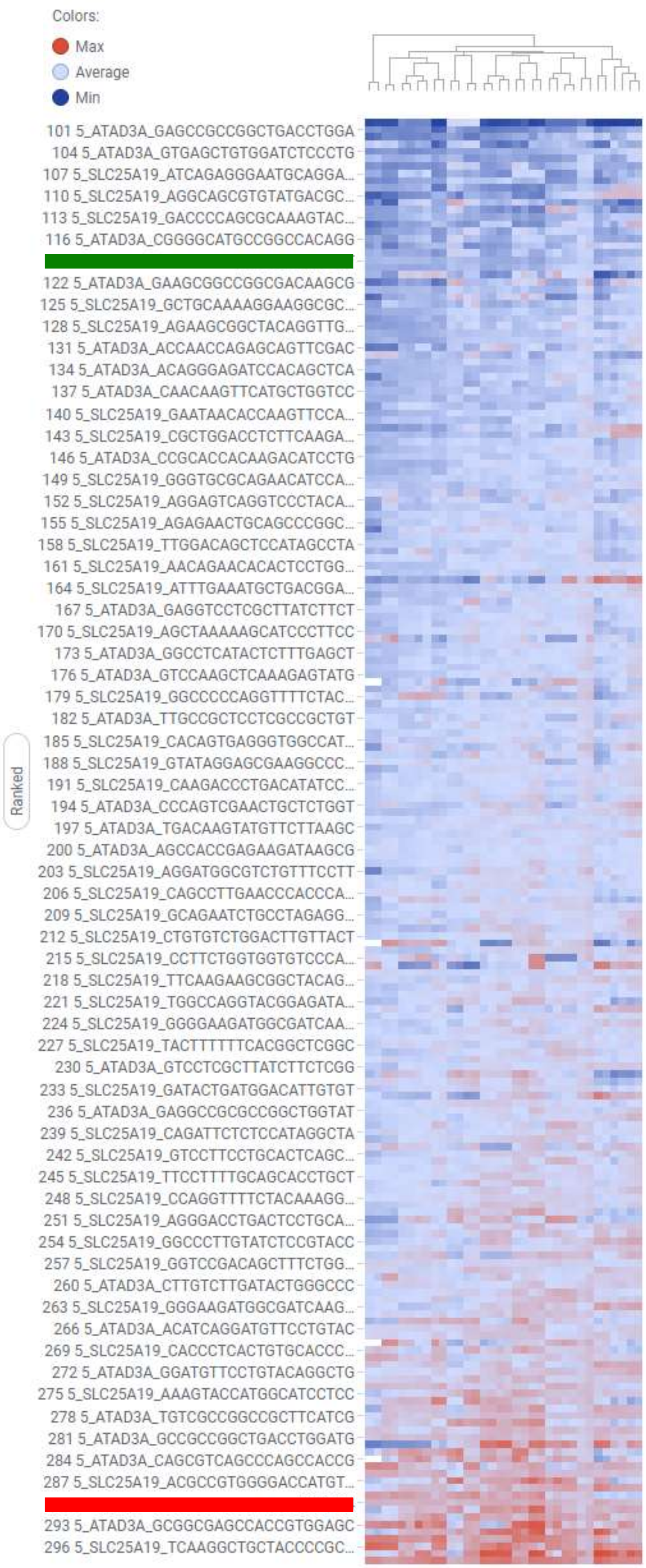


**Figure S10. Low-Feature Count aNNs Rank gRNAs Based on MitoTracker and TMRM Morphologies.** Details on the multi-feature aNNs used to generate preliminary scores to rank SLC25A19 and ATAD3A gRNAs. Each row is a gRNA (only every 3 are labeled), and each column is a model which contains 5, 8, or 15 features within MitoTracker or TMRM wavelengths. The columns are hierarchically clustered in this view. The green bar indicates the Non-Target gRNA and the red, the MFN2.

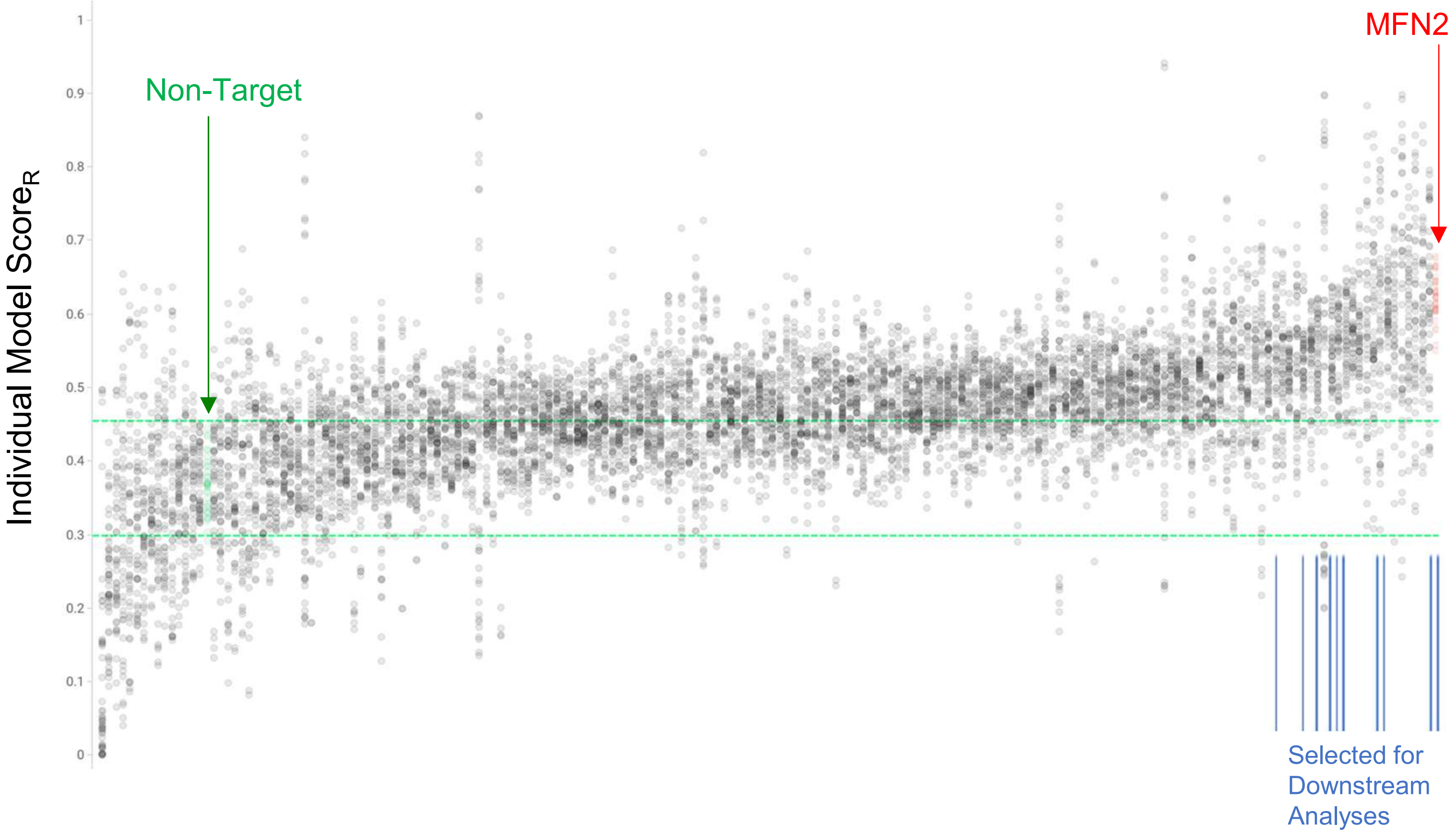


**Figure S11. Individual Model Scores across gRNAs for SLC25A19 and ATAD3A.** Each marker is single-cell image feature data from one of 48 models for each recovered gRNA. The gRNAs are rank-ordered along the X-axis. The position of the Non-Target control and the MFN2 disrupted control are marked. Each of the 48 models are aNNs with three dense layers and either 5, 8, or 15 input features that were solely MitoTracker or TMRM. The position of the gRNAs that were selected for further analysis are demarcated by blue lines.

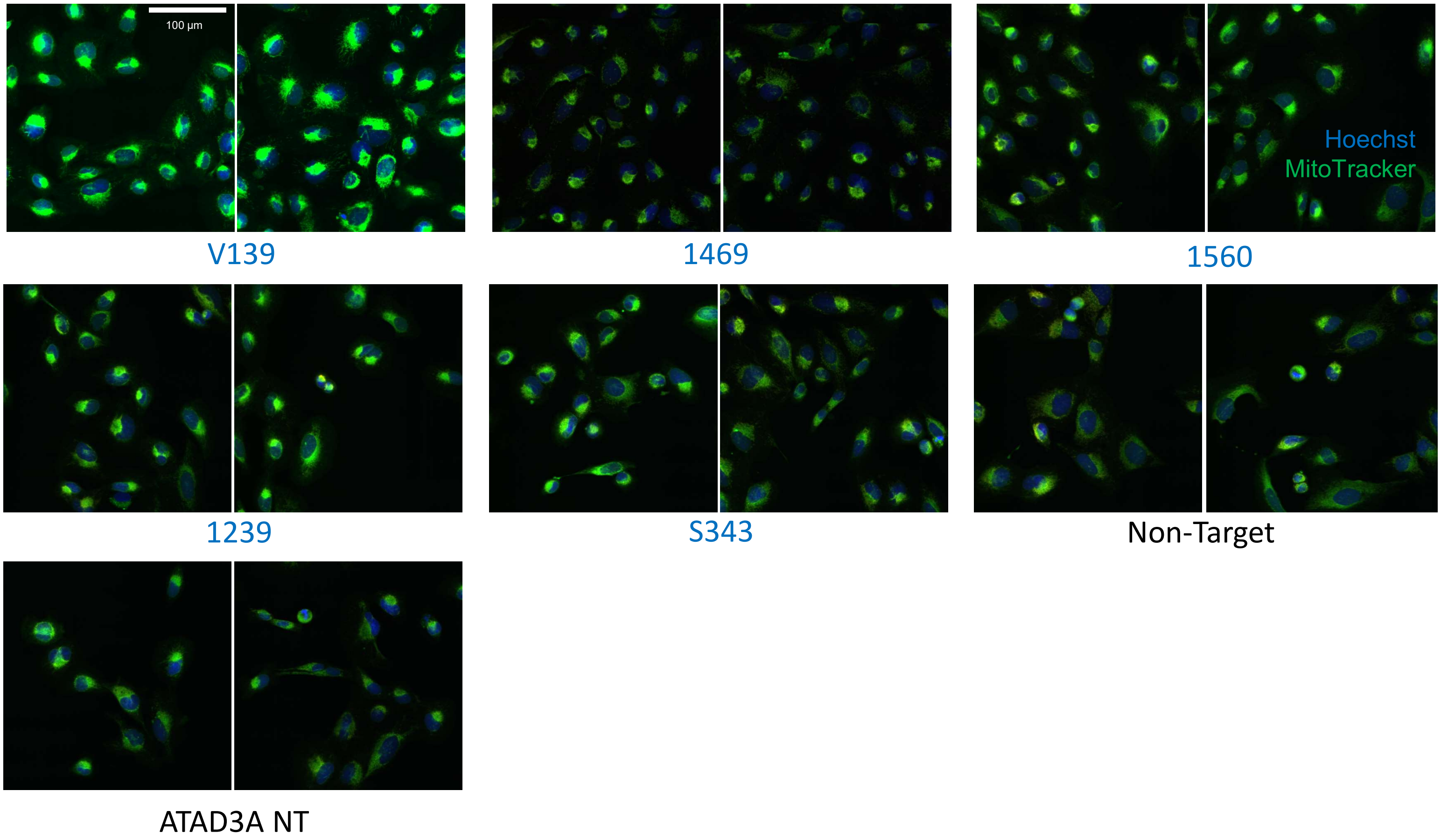


**Figure S12. Cells with Various SLC25A19 and ATAD3A gRNA Edits.** 20x fluorescent images of cells with Hoechst visualized in blue and MitoTracker visualized in green on 96-well plates. Two fields of view are shown for each gRNA. Varying mitochondrial morphologies are visible across the different gRNAs. Labels in blue are for SLC25A19 and mark either the UTR position or the amino acid position.

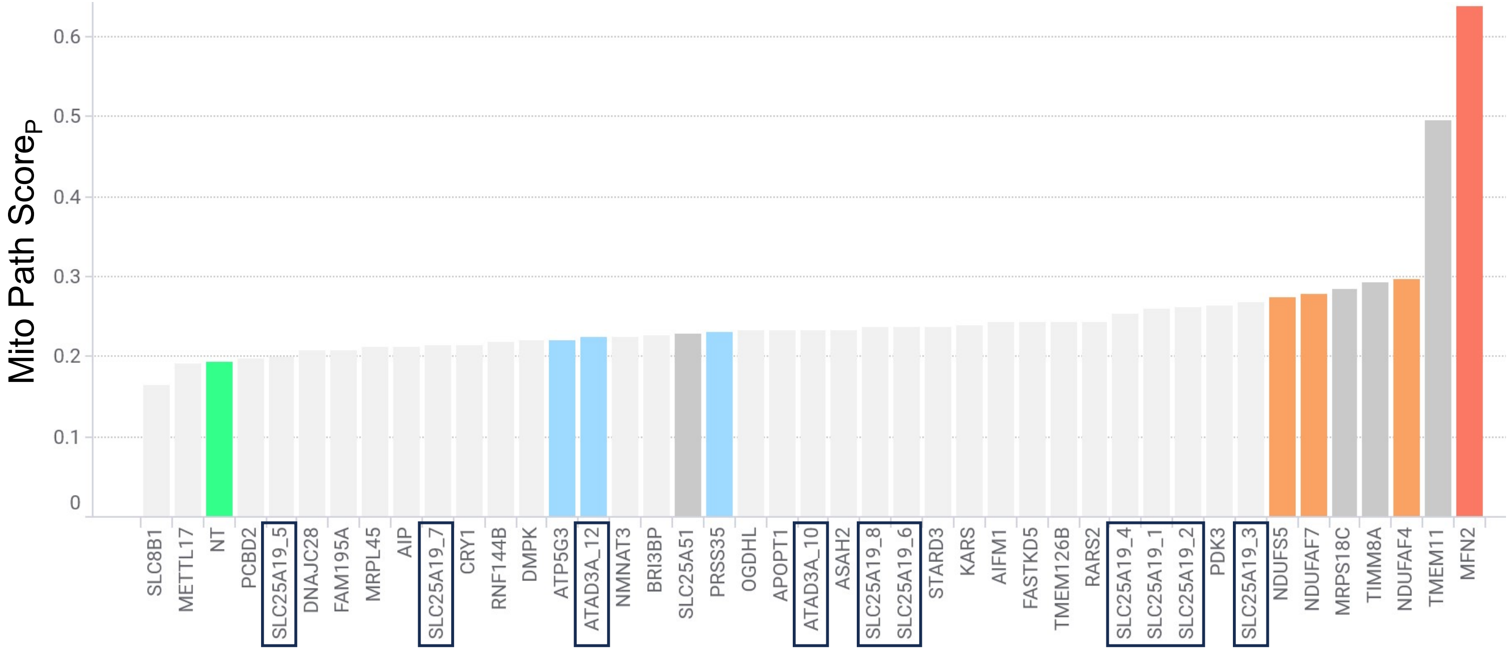


**Figure S13. Comparison of Mito Path Score$_P$ Magnitude Across All Validated gRNAs.** All of the validation results with synthetic gRNAs shown on the same scale. The y-axis shows the Mito Path Score$_P$ , where high numbers are more similar to MFN2 gRNA cells. The x-axis lists the genes which are affected by the various gRNAs. The SLC25A19 gRNAs produce variants which are just below the NDUs in magnitude, while the ATAD3A gRNA is much weaker. SLC25A19 and ATAD3A gRNAs are boxed for easier visibility.

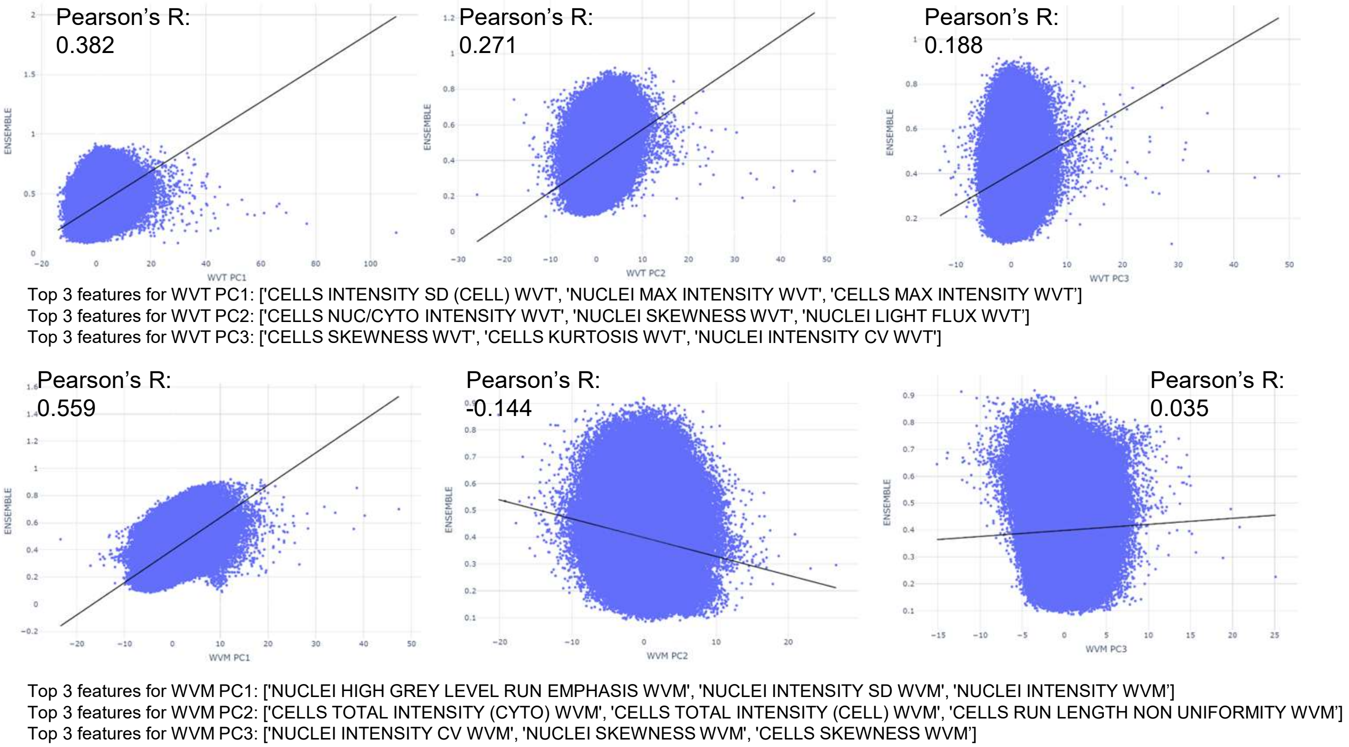


**Figure S14. Combinatorial Principal Component Analysis with Groups of Features per Individual Cell.** Scatter plots with each marker depicting a single cell from the KO gRNA library on microrafts. The y-axis is the Mito Path Score$_R$, and the x-axis is different principal components. The top row of plots are TMRM-feature PCs 1, 2, and 3, and the bottom row are MitoTracker PCs 1,2, and 3. The top three feature loadings are listed below each plot, as well as the Pearson r statistics.

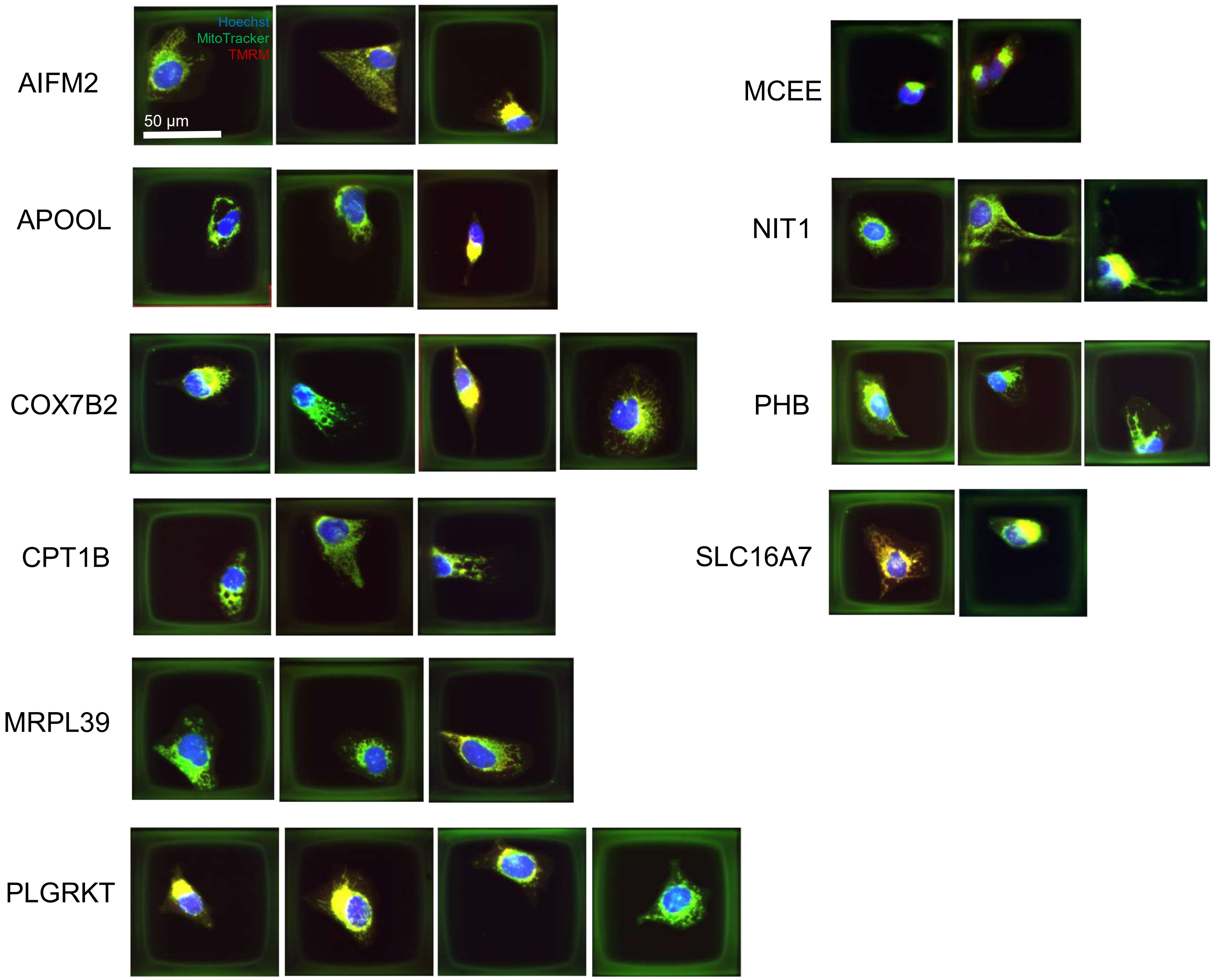


**Figure S15. Microraft Images of Cells with Anomalous gRNAs.** 20x fluorescent images of cells on microrafts of particular recovered gRNAs with the highest Anomaly Score. Hoechst (nuclei) is visualized in blue, MitoTracker is visualized in green, and TMRM is in red (mito+TMRM looks orange or yellow). Various forms of the mitochondria are visible, particularly pertaining to the degree of fragmentation or aggregation.
